# Coordinated immune architecture underlies durable survival in pancreatic cancer

**DOI:** 10.64898/2026.04.29.721612

**Authors:** Nabeel Merali, Sakina Amin, Shihong Wu, Carlos Valverde-Hernandez, Weijia Gao, Tarak Chouari, Chanthirika Ragulan, Hari Ps, Izhar Bagwan, Nariman D. Karanjia, Rajesh Kumar, Rajiv P. Lahiri, Tim D. Pencavel, Esther Platt, Angela Riga, Tim R. Worthington, Sunny Sunshine, Shivan Sivakumar, Anguraj Sadanandam, Timothy A. Rockall, Elisa Giovannetti, Alex Gordon-Weeks, Michael L. Dustin, Nicola E. Annels, Rachael Bashford-Rogers, Adam E. Frampton

## Abstract

Pancreatic ductal adenocarcinoma (PDAC) is characterised by profound immune dysfunction and limited response to immunotherapy. Although tertiary lymphoid structures (TLSs) are associated with improved outcomes, most studies focus on their presence or density, providing limited insight into how their organisation contributes to tumour control. Here, we analyse a unique cohort of treatment-naïve PDAC patients enriched for exceptionally rare long-term survivors (LTS, n = 23; >5-year survival) alongside more common short-term survivors (STS, n = 24), enabling direct interrogation of mechanisms underlying durable tumour control. We develop a multi-scale spatial framework integrating multiplex imaging, computational modelling, and transcriptomics to quantify TLS architecture. By modelling TLSs as higher-order immune assemblies, we define their structural states, spatial organisation, assembly rules and tissue context-dependency within the tumour microenvironment. We show that long-term survival is associated with organised, spatially integrated TLSs exhibiting coordinated B and T cell zoning, whereas short-term survival is characterised by disorganised, Treg-enriched, and spatially isolated TLSs, despite similar immune cell abundance. These architectural differences align with transcriptional programmes: LTS tumours display coordinated lymphoid and NF-κB–driven chemokine signalling, while STS tumours exhibit inflammation uncoupled from immune organisation. Together, these findings demonstrate that durable anti-tumour immunity in PDAC is defined by coordinated transcriptional, cellular, and spatial organisation, rather than immune presence alone. This study provides a blueprint for structure-informed strategies to reprogram the tumour microenvironment and improve outcomes in PDAC.

**Statement of significance:** Integrating multiplex imaging, computational modelling and transcriptomics in treatment-naïve PDAC enriched for exceptional survivors shows that durable tumour control is associated with organised TLS architecture rather than immune abundance alone, providing a framework for immune-architecture-based biomarkers and therapeutic reprogramming.

## Introduction

Pancreatic ductal adenocarcinoma (PDAC) remains one of the deadliest human malignancies, accounting for over 90% of pancreatic cancers and carrying a 5-year survival of just 12% in the United States and 7% in the United Kingdom [1][2]. Fewer than 5% of patients survive beyond a decade. Chemotherapy regimens such as gemcitabine, 5-fluorouracil (5-FU), capecitabine, and FOLFIRINOX offer only modest benefit [3], and while surgical resection provides the only chance of cure, just 20% of patients present with resectable disease [4]. Even novel transformative approaches in other cancers, including immune checkpoint blockade and engineered T-cell therapies [5], have largely failed to shift outcomes in PDAC [6].

Yet within this dismal landscape lies a striking paradox: a small group of patients survive far longer than expected, sometimes for more than eight years [7]. These individuals represent proof that durable control of PDAC is biologically possible. Here we address the critical question of what tumour-intrinsic features, immune dynamics, stromal interactions, or treatment responses enable these exceptional outcomes. Through deciphering the mechanisms that allow certain patients to have exceptional survival, we may uncover actionable vulnerabilities and protective pathways that can be harnessed therapeutically. Long-term survivors (LTS) offer a living blueprint for potential therapeutic success. By understanding the determinants of long-term survivorship, we have the opportunity to transform rare biological success into broadly applicable treatment strategies, shifting PDAC care from prediction to intervention, and from inevitability to possibility.

A defining feature of PDAC is its highly desmoplastic and immunologically complex tumour microenvironment (TME), which acts not merely as a bystander but as a central driver of tumour progression and therapeutic resistance [8]. The dense stromal architecture, coupled with profound immune dysregulation, creates a niche that shields malignant cells from cytotoxic therapies and blunts effective anti-tumour immunity. Within both the peripheral circulation and the TME, immune cell populations exist along a functional spectrum, capable of mediating tumour clearance or, conversely, reinforcing immune escape [9]. PDAC tumorigenesis is tightly linked with the accumulation of immunosuppressive cell subsets. Regulatory T cells (Tregs) [9,10], macrophage M2 [11], myeloid derived suppressor cells (MDSC) [12] and tumour associated macrophages (TAMs) [13], regulatory B cells (Bregs)[14], and dysfunctional T cell populations [15] collectively establish a tolerogenic environment that impairs cytotoxic immune responses. Although the precise role of Tregs in PDAC remains complex and at times context-dependent [16], their presence is associated with poor prognosis [17].

Comparing the immune landscapes of LTS and short-term survivors (STS) offers more than prognostic stratification; it provides a window into the mechanisms that enable durable tumour control in a disease where this is exceptionally rare. Prior studies have contrasted survivors across varying thresholds of survivorship (>8 months, 3 years, 5 years, and 10 years) and identified clinical, genomic, and tumour-intrinsic features associated with outcome [18–20]. However, patients surviving beyond 4.5 years, representing fewer than 2% of all cases [21], remain profoundly underexplored. These exceptional survivors represent a biologically distinct group whose tumours have been effectively controlled for years. Emerging literature underscores the importance of tumour–immune co-evolution in PDAC. Neoantigen editing by the immune system has been implicated in shaping tumour immunogenicity [22], and early-phase studies of personalised mRNA vaccines have demonstrated the capacity to generate neoantigen-specific T cell responses associated with delayed recurrence in resectable disease [23]. Genetic analyses have revealed germline variants in CHEK2, RAD51D, MLH1+ATM enriched in LTSs [24], while spatial transcriptomics has identified immune compositional differences, including increased B cell infiltration in patients surviving beyond three years [25]. Together, these studies suggest that, even in PDAC, meaningful anti-tumour immunity can emerge under specific biological conditions.

A growing body of evidence points to tertiary lymphoid structures (TLS) as a potential anatomical and functional nexus of this immunity. TLS are ectopic lymphoid aggregates that recapitulate key features of secondary lymphoid organs, including B–T cell compartmentalisation, antigen presentation, and local lymphocyte priming [26]. In PDAC, intratumoral TLSs have been associated with favourable prognosis, neoantigen burden, humoral immune responses, and long-term survival, including following neoadjuvant therapy [27–29]. Mechanistically, stromal elements such as cancer-associated fibroblasts may promote lymphoid neogenesis through acting as lymphoid organisers via secretion of CXCL13, CCL19, and CCL21 [27,30], reinforcing the concept that immune architecture is actively organised within the tumour microenvironment.

Despite these advances, critical gaps remain. TLS assessment in PDAC has largely been limited to presence or density, with inconsistent definitions of maturity and limited evaluation of functional activity, such as germinal centre formation. Importantly, features of the spatial organisation likely central to TLS biological function, such as cellular zoning, and microenvironmental context of TLS, have not been systematically integrated with survival outcomes across well-annotated cohorts [31–36]. Without this multi-scale understanding, the field lacks the mechanistic clarity needed to translate TLS biology into robust biomarkers or therapeutic strategies. In this study, we move beyond descriptive TLS quantification to perform a comprehensive spatial and computational characterisation of TLS architecture in PDAC. Focusing on well-characterised exceptional LTSs (>5 years), we systematically define and quantify multi-scale structural, cellular, and spatial features of TLS and evaluate their association with durable survival. By dissecting the immune microarchitecture that underpins rare long-term disease control, this work aims to identify actionable principles that could inform immune reprogramming strategies for the broader population of patients with PDAC.

## Results

### Long-term survival is marked by increased TLS abundance and immune-enriched tumour phenotype

To define the clinical and biological context of exceptional survival in PDAC, we applied an integrated multi-modal framework combining transcriptomics, histopathology, and high-dimensional spatial immune profiling (**Figure 1A-D**). The cohort comprised 47 treatment-naïve resected PDAC patients with tumours located in the head of the pancreas, stratified into long-term survivors (LTS; n=23) and short-term survivors (STS; n=24) (summarised in **Table 1**). The median disease-free survival (DFS) was 80 months in the LTS group vs. 13 months in the STS group (*p* < 0.0001), and median overall-survival (OS) was 90 months versus 23.5 months, respectively (*p* < 0.0001, **Figure 1F**). The baseline age at surgery was comparable between groups (67.5 [49–81] years for LTS vs. 69.1 [53–82] years for STS; *p* = 0.5867, **Table 1**). A history of smoking was more common among STS patients (*p* = 0.0365), with 54% identified as non-smokers, compared with 82% in the LTS group. Adjuvant chemotherapy regimens differed between groups (*p* = 0.0471), with a higher proportion of LTS patients completing their treatment. Tumour size, disease stage (p = 0.0468), and lymph node ratio (p = 0.0276) were higher in the STS group than in the LTS group, consistent with a more aggressive tumour phenotype in STS. Median serum CA 19-9 levels were markedly elevated in STS patients (1278 U/mL [4– 17805]) compared with LTS (193 U/mL [2–1097]; *p* = 0.0014). We next interrogated the immune landscape using nine-colour multiplex immunohistochemistry (DAPI, PanCK, CD19, CD20, CD68, DCLAMP, CD4, CD8, FOXP3) applied across tumour core and invasive regions (**Figure 1A-E**). All tertiary lymphoid structures (TLSs) were manually annotated on digital whole-slide images, enabling quantification of total TLS number and density within tumour regions. Strikingly, long-term survival was associated with a significantly greater number of TLSs (*p* = 0.0370, **Figure 1G**), and TLS density per mm² of tumour was significantly higher in LTS (*p* = 0.0081). These findings confirmed previous observations that increased TLS abundance is a defining spatial feature of durable survival in PDAC [41].

**Figure 1.**
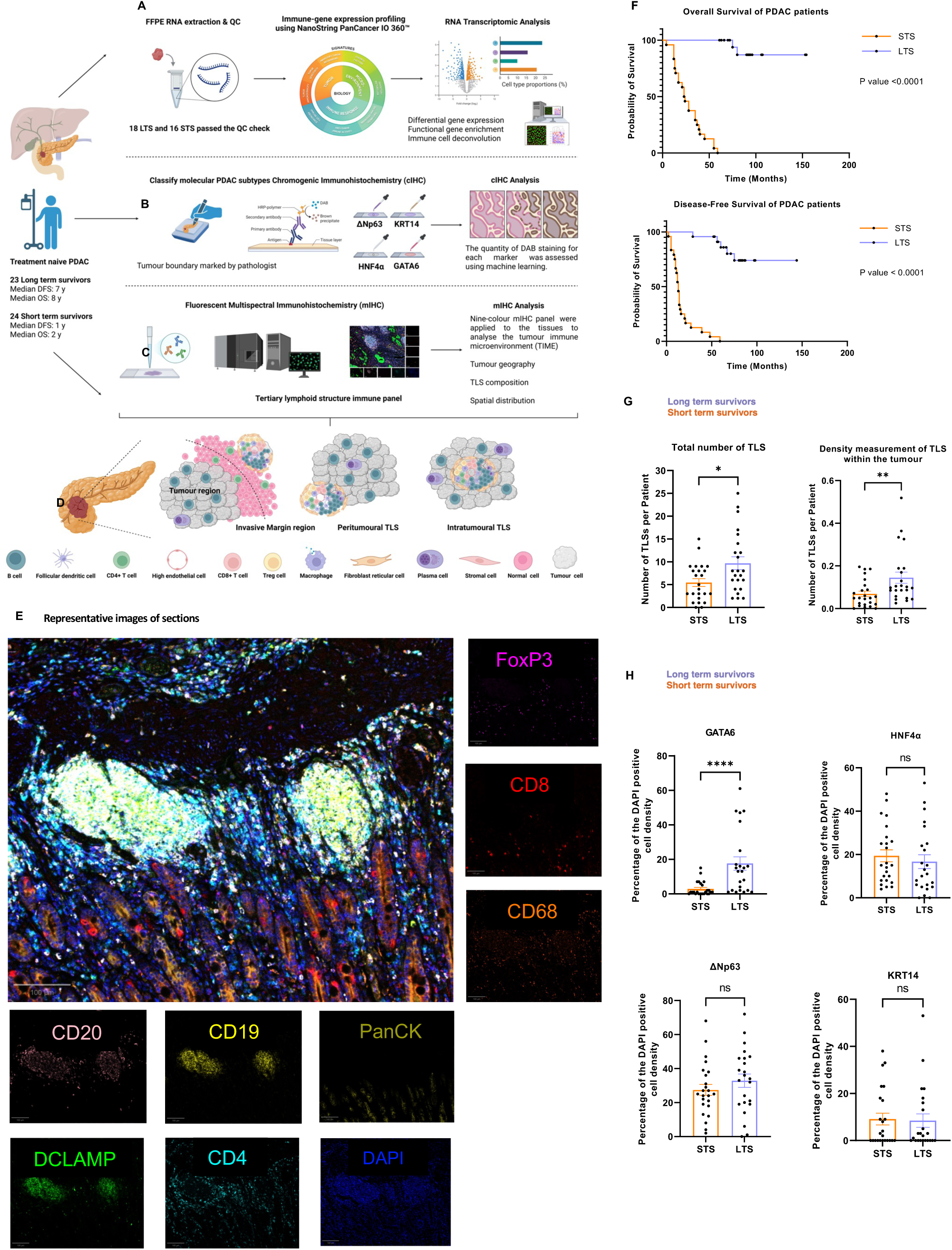
Multi-modal profiling reveals increased TLS burden and immune-enriched phenotype in long-term PDAC survivors. (A) Bulk RNA transcriptomic data extracted from FFPE tumour samples and profiled using the NanoString PanCancer IO 360™ panel from LTS (n=18) and STS (n=16). (B) Chromogenic immunohistochemistry (cIHC) was performed to classify PDAC molecular subtypes (basal-like vs. classical) using established markers (GATA6, HNF4α, ΔNp63 and KRT14) panel from LTS (n=23) and STS (n=24). (C) Fluorescent multispectral immunohistochemistry (mIHC) was applied to multiplexed tumour sections to characterise the composition, spatial distribution, and maturity of tertiary lymphoid structures (TLS). A nine-colour panel enabled simultaneous visualisation of key immune subsets within peritumoural and intratumoural niches. (D) Characterisation of Tertiary Lymphoid Structures (TLS). TLS were identified in three anatomical contexts: the invasive margin, peritumoural regions, and intratumoural compartments. This visual framework highlights the heterogeneity and immune complexity of TLS across distinct tumour microenvironment niches. (E) Representative nine-colour mIHC image of PDAC tissue depicting the spatial organisation of immune and stromal cells within and surrounding a TLS. The composite highlights dense lymphoid aggregation adjacent to tumour epithelium, with clear compartmentalisation characteristic of TLS architecture. mIHC images for CD8 (Opal 480), CD68 (Opal 570), CD4 (Opal 690), DCLAMP (Opal 520), CD19 (Opal 650), FoxP3 (Opal 620), CD20 (Opal 540), PanCK (Opal 780), DAPI and the Composite image (colour legend represented at the bottom right). All images were taken at 20x using the PhenoImager HT (Akoya Biosciences) by our collaborators. Scale bar to 100 μm shown at the left bottom side of each image. (F) Kaplan–Meier overall survival curves for PDAC LTS (n=23) and STS (n=24). Kaplan–Meier survival analysis comparing overall survival and disease-free survival plotted over time (months), with tick marks indicating censored observations. The p-values generated from the log-rank test between the two groups are shown on each plot. Significance thresholds: ns = not significant; *P < 0.05; **P < 0.005; ***P < 0.0005. (G) TLS density (mm^2^) and total number of TLS in LTS (n=23) and STS (n=17). Data are presented as bar graphs showing the mean ± standard error of the mean (SEM), with individual patient values. Statistical comparison between groups was performed using the Mann–Whitney U test. Significance thresholds: ns = not significant; *P < 0.05; **P < 0.005; ***P < 0.0005. (H) Percentage of the DAPI positive cell density values were quantified across tumour sections and compared between LTS (n=23) and STS (n=17). Data are presented as bar graphs showing the mean ± standard error of the mean (SEM), with individual patient values plotted as dots. Statistical comparison between groups was performed using the Mann–Whitney U test. Significance thresholds: ns = not significant; *P < 0.05; **P < 0.005; ***P < 0.0005.

**Table 1.**
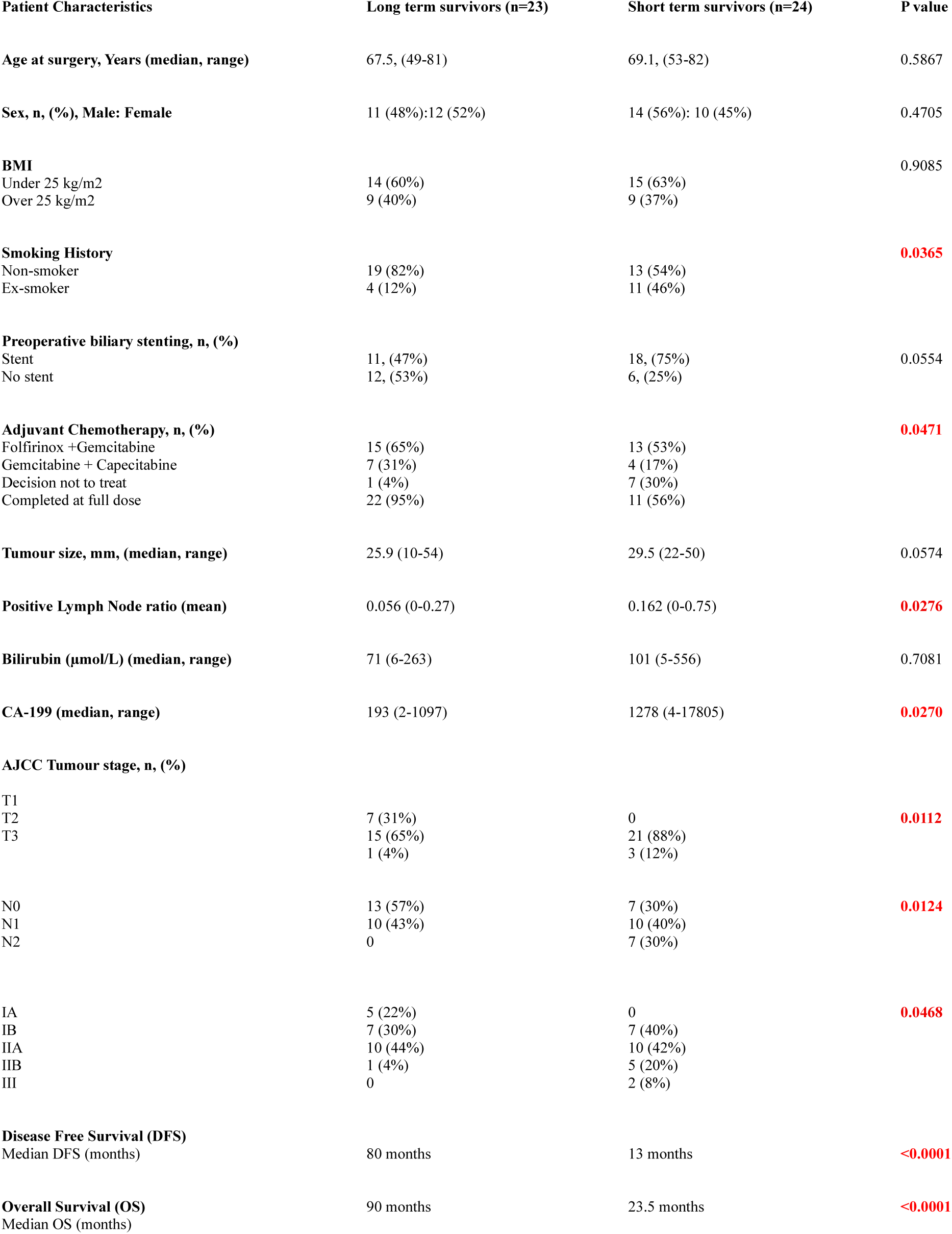
Patient characteristics.

To further contextualise these differences, tumours were molecularly subtyped using chromogenic IHC markers (GATA6, HNF4α, ΔNp63, KRT14) with machine-learning–based quantification (**Figure 1H**). While basal markers (ΔNp63, KRT14 and HNF4α) showed non-significant trends, the classical subtype transcription factor GATA6 was markedly elevated in LTS (*p* < 0.0001). GATA6 has been linked to an immune-enriched tumour phenotype characterised by higher immune infiltration including TLSs and improved outcomes [42–44].

### Global immune cell abundance is similar between groups, but Treg enrichment marks poor outcome

To define regional immune architecture in PDAC, we performed automated single-cell phenotyping and neighbourhood analysis on multiplexed tissue images spanning tumour core and invasive compartments (**Figure 2A**). Cells were assigned to nine major populations (epithelial, non-epithelial, CD4⁺ T cells, CD8⁺ T cells, FOXP3⁺ Tregs, macrophages, dendritic cells, CD19⁺CD20⁺ B cells, and CD19⁺CD20low B cells), and local cellular neighbourhoods were computed within a 30 μm radius. TLS and non-TLS regions were then annotated based on nearest-neighbour structure (**see Methods**), enabling region-specific immune quantification.

**Figure 2.**
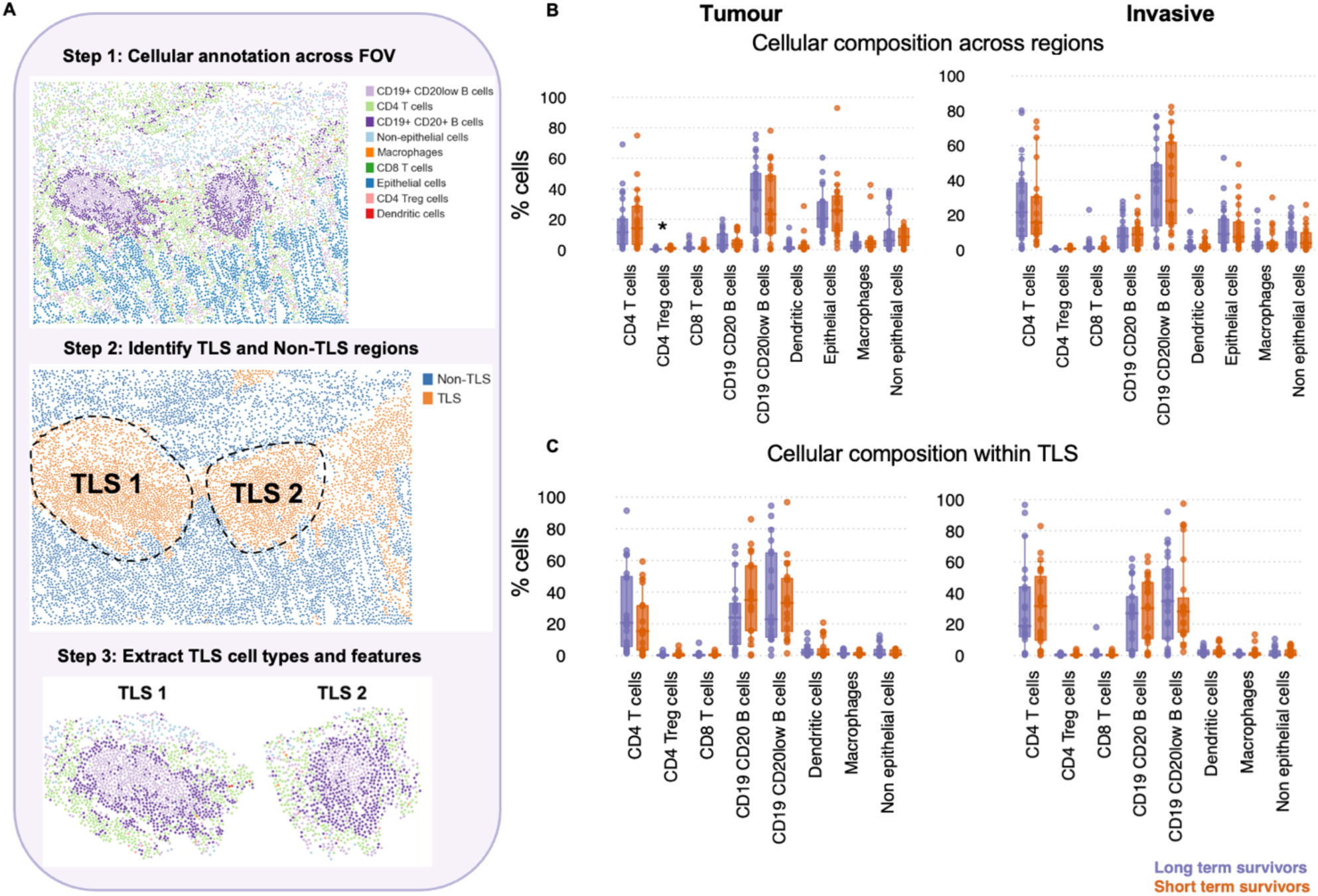
Spatial phenotyping and composition in PDAC reveal Treg enrichment in short-term survivors. (A) Representative field of view (FOV) illustrating the sequential steps for high-dimensional spatial analysis: Step 1 shows automated single-cell phenotyping across the tissue; Step 2 depicts the identification and segmentation of immune-enriched versus non-immune-enriched regions based on nearest-neighbour architecture; Step 3 demonstrates the extraction of specific cellular features and immune subsets within the defined TLS boundaries for multi-scale quantification (B, C) Comparison of the relative abundance of nine major cell populations (including T cell subsets, B cells, myeloid cells, and epithelial/non-epithelial compartments) within the tumour core and invasive margin within the whole FOVs (B) and within TLSs (C). P-values were generated by MANOVA.

As expected, epithelial and stromal compartments dominated the cellular landscape, with considerable inter-patient variability in immune infiltration (**Figure 2B**). Overall, the abundance of CD4⁺ and CD8⁺ T cells, macrophages, dendritic cells, and B cell subsets did not differ significantly between LTS and STS in either tumour core or invasive regions. The distinction between LTS and STS cellular abundances was immunoregulatory: STS tumours harboured a significantly higher proportion of intratumoural FOXP3⁺ Tregs (mean ± SD: 0.91 ± 0.86 vs 0.47 ± 0.50, p = 0.041; **Figure 2B**), consistent with a more immunoregulatory microenvironment in patients with poor clinical outcome. We next examined whether regional differences emerged at the level of spatial organisation by analysing cell-type composition across neighbourhood types. No significant differences were observed within survivor groups, although a trend towards increased B cell representation within tumour regions of LTS was observed (**Figure 2A-C**). Together, these findings indicate that broad immune cell abundance alone does not explain durable survival in PDAC. These results prompted a deeper investigation into higher-order TLS architecture and spatial organisation, features that may better capture the structural immune complexity linked to productive anti-tumour immunity in other solid malignancies such as melanoma and sarcoma [45–49].

### Detailed characterisation of TLS features between long- and short-term survivor groups

To move beyond cell-type abundance and capture higher-order organisation of TLSs, we quantified a comprehensive set of spatial features describing immune architecture at multiple scales (**Figure S2, Figure 3A**). These features included radial distribution metrics (capturing how specific immune populations were positioned relative to TLS cores and margins), geometric descriptors (e.g. axis length), density and distance metrics, and textural/architectural features of TLS organisation (Haralick texture, entropy, and lacunarity) that reflect pattern regularity, complexity, and spatial gaps within TLS regions. To integrate these correlated metrics into biologically interpretable patterns, we performed dimensionality reduction and weighted feature co-expression network analysis (WFCNA), which grouped features into modules representing coordinated spatial programmes (**Figure 3A–B**).

**Figure 3.**
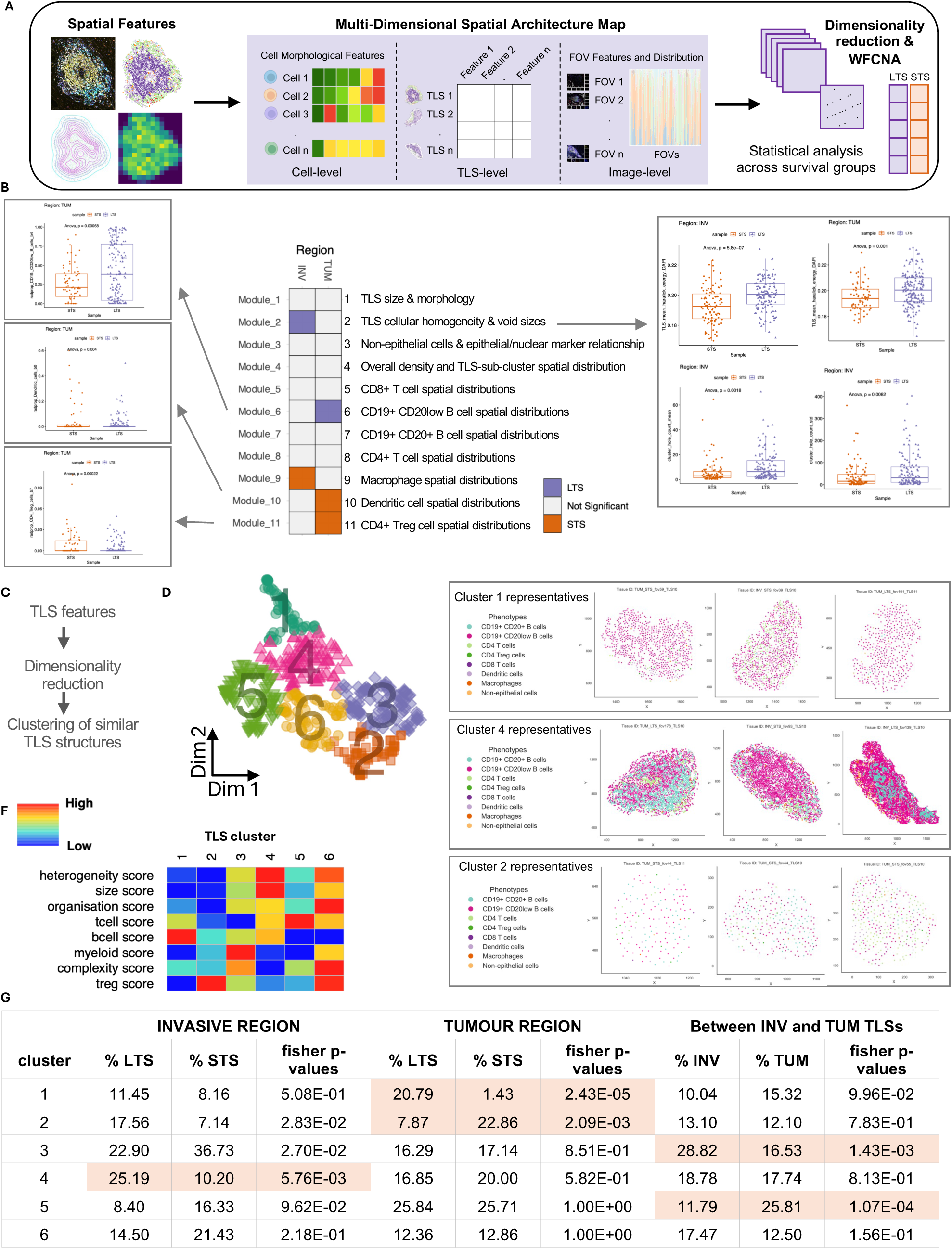
Higher-order spatial organisation and assembly rules of TLSs distinguish long- and short-term survivors in PDAC. (A) Overview of the multi-dimensional spatial architecture framework used to characterise TLS organisation. Spatial features were extracted at the cell, TLS, and field-of-view (FOV) levels and integrated using dimensionality reduction and weighted feature co-expression network analysis (WFCNA) to identify coordinated spatial modules, which were subsequently compared between long and short-term survivors. (B) Heatmap summarising spatial feature modules identified by WFCNA. Module associations with tumour (TUM) and invasive (INV) regions are indicated with representative module scores. (C) Schematic of the pipeline for multi-dimensional TLS characterization, including spatial feature extraction, dimensionality reduction, and unsupervised clustering. (D) UMAP projection of TLS structures across the cohort, coloured by the six identified architectural clusters. (E) Representative spatial maps illustrating the cellular composition of Cluster 1 (B cell-enriched), Cluster 4 (high structural complexity), and Cluster 2 (small, disorganized). Points represent individual cell phenotypes. (F) Heatmap displaying normalized scores (low, blue; high, red) for architectural and immunological metrics across Clusters 1–6. (G) Association of TLS architecture and survival across regions.

A central finding was that TLS architecture and structural integrity differ markedly between survival groups (**Figure 3B**). Haralick energy and entropy metrics showed that TLSs in LTS were more organised, while those in STS were more disordered. Lacunarity analysis further revealed differences in internal void structure, indicating disrupted and less cohesive TLS organisation in STS. Consistently, TLSs in LTS were larger and more spatially developed, with increased convex area, whereas STS TLSs were smaller and less structurally coherent (**Figure S3**). The multiplex panel was designed to resolve major immune and epithelial compartments, enabling high-resolution characterisation of lymphoid organisation within TLSs. Accordingly, lacunarity, entropy, and related texture metrics capture the spatial complexity of TLS architecture as a whole reflecting variation in immune-cell packing density, intercellular spacing, and zonal boundaries rather than the distribution of any single cell population.

At the cellular level, spatial positioning was the key discriminator. **Figure S5** illustrates the spatial identification of TLS on IHC images, along with their segmentation and cellular composition within tumour tissue, including delineation of TLS boundary cells. These analyses highlight that TLSs are not uniform structures but instead exhibit diverse cellular organisation and immune phenotypes across samples and within the same patient. Module 6 showed that CD19⁺CD20 low B cells were preferentially localised to inner TLS regions in LTS, consistent with organised B cell zoning and functional lymphoid architecture [47,50–52]. This pattern was absent in STS. In contrast, STS TLSs were enriched for immunoregulatory features, with increased FOXP3⁺ Treg localisation (Module 9) across tumour region despite similar overall abundance cellular abundance (**Figure 2C**). Modules 10 and 11 further revealed altered macrophage and dendritic cell positioning in STS, indicating disrupted antigen presentation and immune coordination. Together, these results show that while overall TLS cellular composition is similar, their architecture, organisation, and immune functional states diverge sharply. STS is characterised by disordered, immunosuppressive TLSs, whereas LTS exhibits larger, well-organised TLSs with coordinated immune zoning, highlighting that higher-order structure, rather than cell presence alone, underpins durable anti-tumour immunity in PDAC.

### Distinct TLS architectural states associate with survival in PDAC

To systematically capture heterogeneity in TLS organisation, we performed dimensionality reduction followed by unsupervised clustering of multi-scale spatial features, identifying six reproducible TLS states spanning a continuum from small, poorly organised aggregates to large, architecturally complex lymphoid structures (**Figure 3C–F**). These states reflected clear structural and immunological differences. Clusters 1, 2 and 4 were significantly different between PDAC LTS and STS groups. **Cluster 4** represented the most developed TLSs, and were the largest, most highly organised and heterogeneous, consistent with expanded lymphoid structures containing coordinated immune compartments. **Cluster 1** showed strong B cell enrichment with intermediate structural complexity. In contrast, **Cluster 2** comprised the smallest and least organised TLSs, with high Treg representation, consistent with immature or dysfunctional lymphoid assemblies. The remaining clusters captured intermediate stages, suggesting a spectrum of TLS maturation. **Supplementary Figure S4** shows the other cluster (3,5 and 6) representatives.

TLS state distribution also varied by anatomical context (**Figure 3G**). While all clusters were observed in both tumour core (TUM) and invasive margin (INV), INV regions were enriched for myeloid-associated TLS states (Cluster 3), whereas TUM regions more frequently exhibited T cell–enriched states (Cluster 5) (*p* < 0.01). This highlights the need to consider spatial context when interpreting TLS biology.

TLS architectural states were also strongly associated with survival (**Figure 3G**). In invasive regions, Cluster 4 TLSs, representing the largest and most organised structures, were significantly enriched in LTSs compared with STSs (25.2% vs. 10.2%, *p* = 0.0058). In tumour cores, Cluster 1 TLSs (B cell–rich, organised structures) were markedly enriched in LTS (20.1% vs. 1.43%, *p*= 2.43x10⁻⁵), whereas Cluster 2 TLSs (small, disorganised, Treg-enriched) were more frequent in STS (22.86% vs. 7.8%, *p* = 2.09x10⁻³).

Notably, TLS states were not randomly distributed within tumours. Patients tended to harbour consistent TLS architectures across their tumours (Chi-squared test *p*-value = 1.10x10^-4^, Adjusted Rand Index (ARI) with Permutation Test *p*-value = 6.80x10^-3^), suggesting that TLS organization reflects a tumour-intrinsic immune programme rather than local stochastic variation.

Together, these findings demonstrate that TLSs in PDAC exist along a spectrum of structural states defined by size, organization, and immune composition, and that these states are tightly linked to patient and clinical outcome. Favourable survival is associated with large, integrated, and B cell–organised TLSs, whereas poor outcome is characterised by smaller, disordered, and immunoregulatory TLS configurations, supporting a model in which effective anti-tumour immunity depends on the formation of structurally mature and coordinated lymphoid niches.

### Higher-order organisation of TLSs differs between long- and short-term survivors

To investigate how TLSs are organised beyond individual cell distributions, we adapted the Tissue Schematics framework [53]. Rather than treating each TLS as a single, uniform cluster, we decomposed it into smaller regions, or “neighbourhoods,” each defined by distinct immune cell compositions. We then identified recurring patterns of how these neighbourhoods are arranged, referred to as “*motifs*”, and analysed how these motifs connect to form larger, structured immune assemblies, or “*rules*”. By mapping these relationships, we were able to characterise and quantify how TLSs are organised within the tissue (**Figure 4A**). This approach moves beyond simply identifying which cells are present, enabling us to understand how immune cells are spatially arranged and interact to form functional structures.

**Figure 4.**
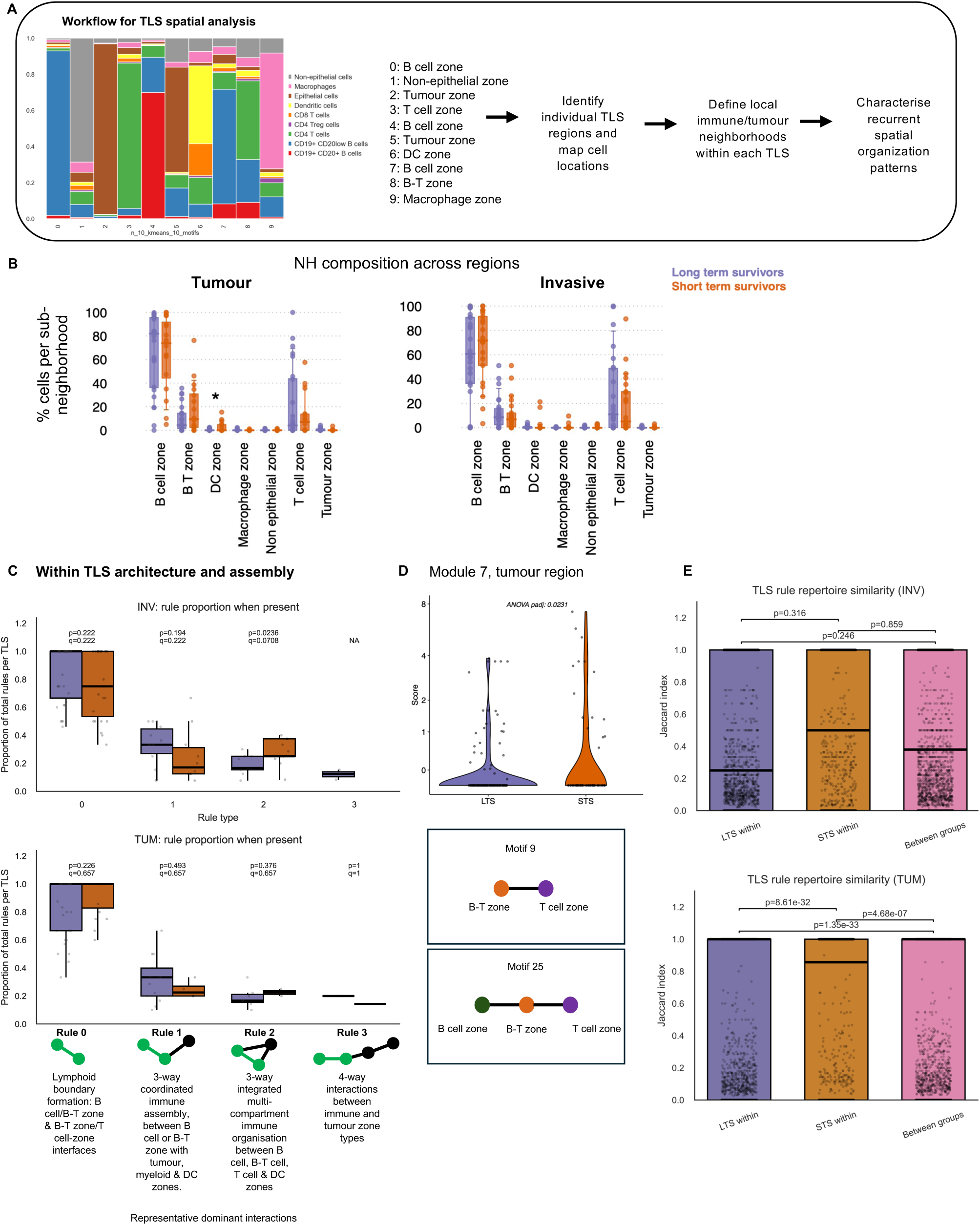
Higher-order TLS architecture and assembly differ between long- and short-term survivors in PDAC. (A) Schematic of the TLS analysis workflow. Individual TLSs were segmented, local cellular neighbourhoods were defined, and recurrent higher-order motifs and assembly rules were derived. Neighbourhoods were annotated as B-cell, B–T, DC, macrophage, non-epithelial, T-cell, or tumour zones. (B) Neighbourhood composition across tumour and invasive TLSs in long-term survivors (LTS, blue) and short-term survivors (STS, orange). (C) Within-TLS rule usage in invasive (top) and tumour (bottom) regions. Boxplots show the proportion of total rules per TLS attributable to each rule family, restricted to TLSs in which that rule was present. Rule families are shown with representative biological interpretations: Rule 0, pairwise lymphoid interfaces; Rule 1, coordinated 3-compartment immune assembly; Rule 2, integrated multicompartment immune organization; Rule 3, rare 4-compartment immune–tumour/stromal interactions. P and FDR-adjusted q values are shown. (D) Tumour-region motif module analysis. Schematics show the dominant contributing motifs (E) TLS rule repertoire similarity within and between survival groups. Pairwise Jaccard similarity of exact rule repertoires was calculated for invasive (top) and tumour (bottom) TLSs.

Each TLS was therefore broken down into smaller sub-regions based on the types of cells present and how they were spatially grouped. This revealed neighbourhoods enriched for key immune populations, including B cells, T cells, dendritic cells, and macrophages, as well as mixed regions where different cell types, such as B and T cells, co-localised, reflecting zones of immune interaction. Across both tumour core and invasive regions, the overall distribution of these neighbourhood types was largely similar between long- and short-term survivors, indicating that differences in outcome were not driven by which immune compartments were present. The primary neighbourhood compositional difference was an increased representation of dendritic cell (DC)–enriched neighbourhoods within tumour-associated TLSs in STSs (**Figure 4B**). Importantly, this suggests that DC abundance alone is not associated with outcome, but their spatial organisation is. In STS tumours, DCs appear more spatially segregated, with reduced co-localization with key T cell subsets, potentially limiting effective antigen presentation to B cells and T cell priming. This altered DC zoning is consistent with the broader disruption of immune coordination observed in STS TLSs and may contribute to impaired anti-tumour responses. While increased infiltration of certain mature DCs can be associated with better survival, an increased level of immature, immune-tolerant DCs has been shown to result in shorter survival and poor prognosis [54]. Mature dendritic cells enriched in immunoregulatory molecules (mregDCs), also known as DC3s or LAMP3+ DCs, are a distinct subset that accumulate in the TME of PDAC. While they possess the potential for high antigen presentation, in the context of PDAC, they function primarily as immunosuppressive cells that limit T cell activation, contributing to the cancer’s high resistance to immunotherapy [55,56].

However, important differences emerged when we examined how these neighbourhoods were organised relative to each other. Using a rule-based analysis, we identified recurring patterns in how different neighbourhood types were arranged and connected within TLSs (**Figure 4C**). These results showed that survival differences were associated with how immune cells were organised into larger, coordinated structures. Across both tumour and invasive regions, the most common patterns reflected simple interactions between lymphoid compartments, particularly boundaries between B cells, B–T mixed zones, and T-cell regions. More complex patterns captured coordinated organisation across multiple compartments, extending lymphoid zones to include DC- or tumour-associated regions. When all TLSs were analysed together, one specific pattern (rule 2) was significantly more frequent in STS than in LTS (*p* = 0.00563; **Figure S6A**). When analyzed by location, this difference was driven specifically by TLSs at the invasive margin, where rule 2 was enriched in STS (*p* = 0.0236; **Figure 4C**), with no significant difference in tumour-core TLSs. Together, these findings suggest that survival differences are not due to the presence or absence of particular immune structures, but rather subtle changes in how local immune interactions are combined into larger TLS architectures, particularly at the invasive edge. Other complex interactions were observed beyond the 4 rules, but were much less common and not statistically testable.

To resolve these coordinated structures into finer patterns, we calculated per-TLS motif proportions and applied VDJ-REMIX to the tumour-region motif matrix to identify correlated motif modules and derive a module score summarising each co-occurring motif programme (**Figure 4D, Figure S6B**). Across all modules, Module 7 scores were higher in STS tumour TLSs (adjusted ANOVA *p* = 0.0236). This module was anchored by motif 9, representing a B–T-zone/T-cell-zone interface, and motif 25, representing a three-compartment B-cell/B–T-zone/T-cell assembly, with related B-cell/B–T and tumour-adjacent variants also contributing. Because this analysis was performed on motif proportions rather than raw counts, this signal reflects relative over-representation of these local motifs rather than simply larger or more motif-dense TLSs.

Finally, we assessed the consistency of higher-order TLS organisation across patients by comparing rule usage similarity using the Jaccard index (**Figure 4E**). In tumour-associated TLSs, LTS exhibited highly consistent rule repertoires (median similarity = 1), indicating that similar architectural patterns were reproducibly assembled across these tumours. In contrast, STS showed significantly lower consistency (*p* < 1x10⁻¹⁰), reflecting greater variability and less coordinated organisation. This difference was specific to the tumour core. In invasive regions, rule-repertoire similarity was lower overall and did not differ significantly between survival groups, suggesting a more heterogeneous and context-dependent organization at the tumour boundary.

Together, these findings indicate that durable survival is associated with a coherent and reproducible tumour-associated TLS assembly programme, whereas short-term survival is characterised by heterogeneous and less coordinated architectures. Notably, STS TLSs were enriched for repeated local B-T cell and T cell interface motifs, but these were embedded within a less integrated overall structure. These results highlight that effective anti-tumour immunity is linked to consistent, coordinated higher-order immune architecture within the tumour, rather than isolated or variable local interactions, whereas TLSs at the tumour margin were more heterogeneous.

Together, this shows that while TLSs in PDAC may contain similar immune components, their higher-order organisation differs between patients, and this organisation may be a key determinant of effective intra-tumoural anti-tumour immunity.

### Coordinated immune activation drives TLS maturation in LTS, while STS exhibit dysregulated inflammation

To complement spatial analyses of TLS architecture, we interrogated tumour immune transcriptional programmes using the NanoString PanCancer IO 360™ panel. Differential expression analysis identified a focused set of significantly altered genes (BY-adjusted *p* < 0.05), with the majority upregulated in long-term survivors (LTS) (**Figure 5A**). In total, 285 differentially expressed genes were identified (**Supplementary Materials, Table S3**).

**Figure 5.**
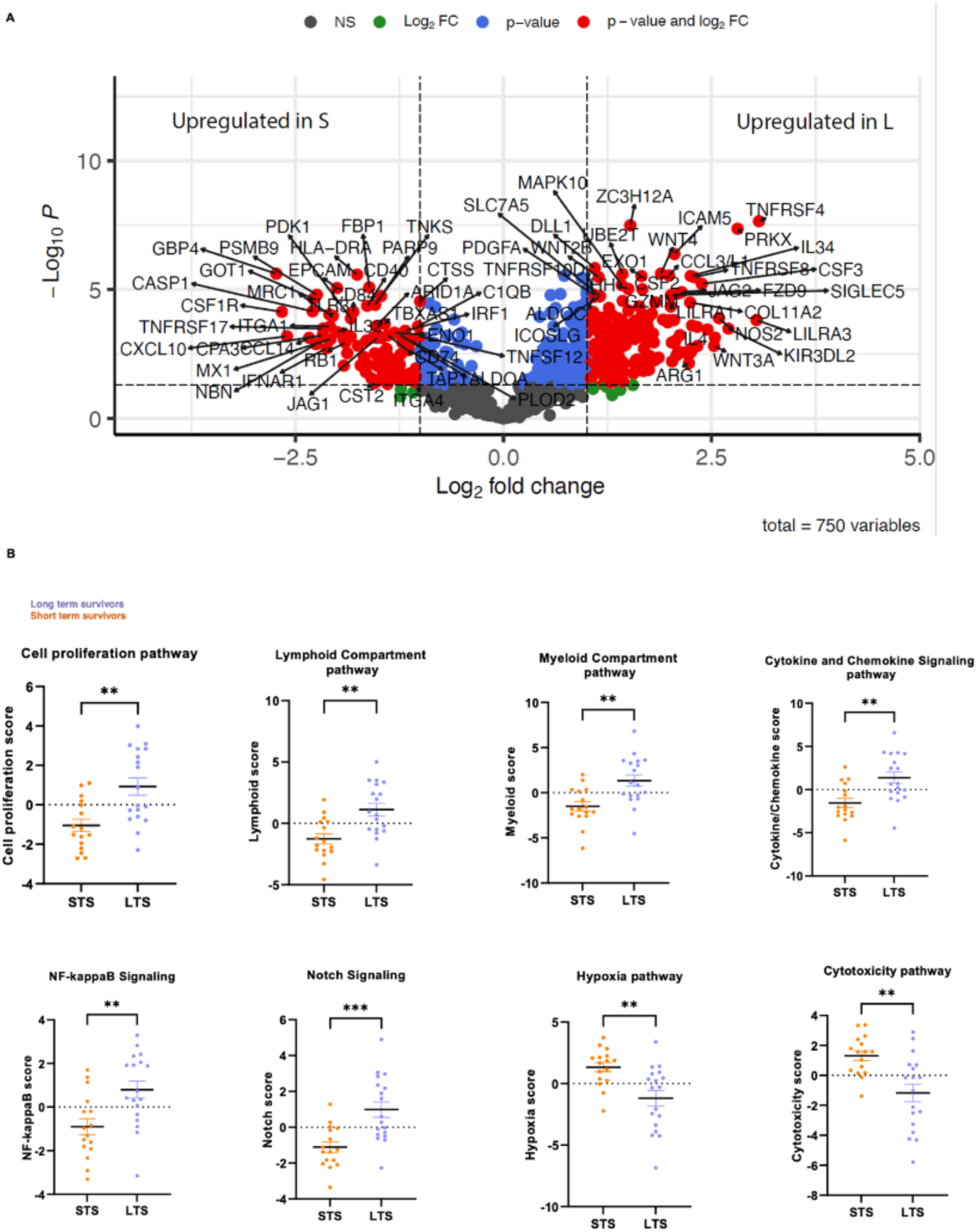
Divergent immune and transcriptional programmes distinguish long- and short-term PDAC survivors. *(A)* Differential gene expression between LTS (n=18) and STS (n=16). Volcano plot illustrating differential mRNA expression in tumour samples from LTS compared with STS patients. The x-axis represents the log₂(fold change) in gene expression and the y-axis displays the –log₁₀(adjusted P value). Each point corresponds to a single mRNA gene transcript. Multiple significance thresholds are indicated: solid line: adjusted P < 0.01, long-dash line: adjusted P < 0.05, short-dash line: adjusted P < 0.10, dotted line: adjusted P < 0.50. *(B)* Significant Differential gene set pathway activation between LTS (n=18) and STS (n=16). Scatter plot data are shown as individual samples with mean ± SEM. Statistical comparison between groups was performed using the Mann–Whitney U test. Significance thresholds: ns = not significant; *P < 0.05; **P < 0.005; ***P < 0.0005.

Among the most significantly upregulated genes in LTS were Immune activation (TNFRSF4, PRF1, GZMM, TBX21), chemokines & cytokines (CCL3, CCL7, IL4, IL12RB2), Immune cell recruitment & signalling (XCL1) and tumour antigen recognition (MAGE family). We have shown that LTS have a highly active, cytotoxic, immune-engaged tumour environment. LTS tumours also exhibited increased expression of SIGLEC5, PDGFA, and JAG2, reflecting immune regulatory, stromal, and Notch signalling pathways, respectively. Significantly elevated WNT2B and ALDOC, indicated activation of developmental and metabolic programmes. Together, these findings suggest that LTS tumours engage multiple coordinated signalling axes spanning immune activation, stromal interaction, and tissue remodelling.

To capture broader biological programmes, we next analysed pathway-level activity using predefined NanoString gene signatures (**Figure 5B**). This revealed a clear divergence in immune states between survival groups. Long-term survivor (LTS) tumours showed increased activity of pathways linked to lymphoid organisation, cytokine and chemokine signalling, NF-κB signalling, and myeloid and lymphoid compartments (*p* < 0.01), consistent with an immune-enriched and functionally active microenvironment. These pathways align with established mechanisms of TLS formation, where NF-κB integrates inflammatory signals such as TNF and lymphotoxin to drive chemokine production (e.g. CCL19, CCL21, CXCL12, CXCL13) and coordinate lymphocyte recruitment and organisation into structured immune niches [57–61]. Accordingly, enrichment of these programmes in LTS suggests effective immune cell recruitment and organisation into functional TLSs, in line with prior studies linking TLS-associated chemokine signatures to improved outcomes across cancers, including PDAC. In contrast, short-term survivor (STS) tumours lacked enrichment of these coordinated pathways despite evidence of inflammation at the gene level, indicating a failure to translate inflammatory signals into organised immune structures. Gene signatures for the detection of tertiary lymphoid structures, derived from transcriptomic analyses of human cancers, are presented in **Supplementary Materials, Table S4**. From these, 11 key TLS-associated signatures were identified.

In contrast, the significantly upregulated genes in STS were myeloid suppression signatures (CSF1R, MRC1, LYZ, S100A9) reflective of a M2-like pro-tumoral macrophage environment [62] with a dense fibrotic matrix with desmoplasia (FAP, COL5A1, COL6A3, ITGA1) [63]. Hypoxia and metabolic stress (HIF1A, PDK1) genes showed that the STS tumours were associated with aggressive disease. STS tumours demonstrated enrichment of pathways associated with inflammatory responses, stress signalling, and dysregulated immune activation, including antigen processing and response-to-stimulus pathways. Despite evidence of inflammatory signalling, these programmes appeared uncoupled from effective immune organisation, suggesting a non-productive or chronically activated immune state. A significantly increased cytotoxicity in the STS tumours may be associated with an exhausted cytotoxic cell display without effective killing targets and a hypoxic/immune cold microenvironment. For example, key immune cell surface and interaction genes were upregulated in STS tumours (CD40, CD84, CD48, CD5), however were likely exhausted. In STS tumours, GBP4 and related interferon-inducible GTPases such as GBP1 and GBP2 were up-regulated, consistent with an interferon-driven, immune-infiltrated tumour microenvironment [64]. High GBP4 expression in pancreatic cancer has been associated with T-cell exhaustion and poorer survival, suggesting that although immune cells are present, they are functionally impaired [65]. Kisling et al, identified a HOXA10-driven 5-gene signature enriched in STS, linked to pathways of proliferation, immune evasion and immunosuppression rather than effective cytotoxicity [66]. Similarly, CTSS gene was up-regulated in STS tumours. CTSS acts as a tumour promoter aiding invasion, angiogenesis and immune evasion leading to matrix remodelling and metastasis [67].

Importantly, these transcriptional patterns mirror the spatial phenotypes observed in TLS architecture and mIHC, thereby validating our findings across complementary modalities. The LTS group showed a greater abundance of immune cell populations, including B cells, neutrophils, NK CD56^dim^ cells, and CD8⁺ T cells (**Figure S9**). In contrast, STS tumours were enriched in macrophages, potentially reflecting an increased presence of pro-tumorigenic M2-like subsets. Gene Ontology (GO) and pathway enrichment analysis in LTS further demonstrated significant overrepresentation of immune-related pathways, particularly those involved in leukocyte activation, T cell signalling, and cytokine-mediated immune responses.

In LTS, elevated lymphoid and cytokine signalling aligns with the presence of larger, structurally organised TLSs with coordinated immune zoning. In contrast, STS tumours exhibit transcriptional features consistent with inflammation coupled to immune suppression and stromal dysregulation, corresponding to disorganised, Treg-enriched TLSs observed spatially. Taken together, these results demonstrate that survival-associated differences in PDAC are not defined by the presence of immune activation alone, but by its organisation and functional orientation. LTS tumours display a coordinated immune programme that supports TLS maturation and effective anti-tumour immunity, whereas STS tumours exhibit a dysregulated inflammatory state that fails to translate into structured, functional immune architecture.

Together, these findings show that LTS tumours recapitulate canonical TLS-forming programmes, whereas STS tumours exhibit a disconnect between inflammation and immune organisation, highlighting that effective anti-tumour immunity depends on structured, TLS-driven responses rather than inflammation alone.

## Discussion

PDAC remains one of the most immunologically refractory malignancies, characterised by profound immune suppression and limited response to immunotherapy. While TLSs have emerged as favourable prognostic features across multiple cancers studies [68–70] in PDAC have largely focused on their presence or density [33,71,72], leaving their functional organisation poorly understood. Here, by integrating spatial profiling, computational modelling, and transcriptomics, we show that long-term survival is defined not simply by TLS abundance, but by their higher-order architecture, organisation, and integration within the TME. Specifically, structured compartmentalisation, coordinated neighbourhood assembly, and organised B-cell zoning distinguish tumours in survivors. These findings support a model in which effective anti-tumour immunity in PDAC is defined less by immune cell quantity and more by the organisation and integration of immune structures within the TME.

A central insight emerging from this study is that differences in clinical outcome were not explained by broad changes in immune cell abundance, but rather by differences in the spatial organisation of immune populations within the TME. However, STS tumours exhibited a significantly higher proportion of FOXP3⁺ Tregs within the tumour region, reflecting an immunosuppressive tumour microenvironment that suppresses effective cytotoxic immunity (**Figure S7**). Similarly, Pu et al. found high intra-tumoural IL-33⁺ FOXP3⁺ Tregs within PDAC tissues and was linked to poorer OS and DFS with an exhausted CD8⁺ T-cell phenotype [73]. Liu et al. showed that patients with PDAC after surgery that had FOXP3⁺ Tregs enrichment within intratumoral regions led to a reduced DFS and the multivariate analysis confirmed Tregs as a negative prognostic factor [74]. Kiryu et al. noted that even when total TILs were high in PDAC, elevated FOXP3⁺ cells still predicted poorer outcomes supporting the idea of an immunoregulatory microenvironment overriding immune activation [75].

Our higher-order spatial analyses reveal that the organisation of immune cells within TLSs is a key determinant of survival. LTS tumours exhibited structured TLSs with central B cell localisation and coordinated integration with surrounding immune neighbourhoods, including B and T cell zones. In contrast, STS TLSs exhibited less coherent zoning and more fragmented compartmental organisation, representative images are shown in **Figure S8**. Notably, we also observed differences in DC spatial organisation, with tumour-associated TLSs showing distinct DC-enriched zones. Given the central role of DCs in antigen presentation and T cell priming, their localisation within TLSs is likely to be functionally important. In LTS, the organisation of DCs within structured TLSs may facilitate efficient antigen presentation and coordinated activation of adaptive immune responses. Conversely, the reduced positioning of DCs in STS TLSs suggests impaired antigen presentation capacity, potentially contributing to ineffective T cell activation despite the presence of immune cells. These findings highlight that spatial arrangement of antigen-presenting cells, alongside lymphoid organisation, is critical for generating functional anti-tumour immunity. These observations extend prior work demonstrating structured compartmentalisation within PDAC TLSs [75] and reinforce the concept that TLS functionality depends on coordinated cellular positioning, not simply immune cell presence. Sidiropoulos et al. showed that responders to therapy exhibit TLS-associated B-cell maturation and functional immune activation [76]. However, most prior studies have focused on TLS presence or proximity metrics [68–72]. In contrast, our work introduces a multi-scale framework that quantifies TLS architecture, structural complexity, and spatial integration, revealing that favourable outcome is associated with TLSs embedded within coordinated immune assemblies rather than existing as isolated structures.

A further key advance is the integration of spatial architecture with transcriptional profiling. LTS tumours exhibited enrichment of lymphoid organisation, cytokine signalling, and NF-κB–driven pathways, consistent with canonical TLS formation programmes and effective immune coordination. We interrogated published TLS gene signatures [77] and identified a set of 11 key genes (CCL3/L1, CCL4, CCL8, CXCL10, TNFRSF17, ICOSLG, TIGIT, PDCD1, CD40, CD38, and CD5) associated with TLS formation. These genes are derived from previously reported TLS-associated transcriptional programmes. For example, Coppola et al. described a 12-chemokine gene signature in colorectal cancer that encompassed myeloid, T cell, and B cell attractants and showed strong correlation with TLS presence by IHC [78]. Similarly, Gu-Trantien et al. identified an 8-gene T follicular helper (Tfh) cell signature in breast cancer linked to B cell recruitment, Tfh cell function, and the formation of mature germinal centre-like TLS structures [79]. Additional TLS-associated signatures have been described across tumour types, including a T helper 1 (Th1)/B cell signature in gastric cancer [80] and a plasma cell signature in ovarian cancer [81], both of which are associated with TLS-dependent processes such as lymphocyte proliferation, dendritic cell differentiation, and adaptive immune activation. In contrast, STS tumours showed inflammatory and immunoregulatory signatures that were uncoupled from spatial organisation, reflecting a non-productive immune state. These findings highlight that inflammation alone is insufficient, and immune responses must be spatially organised to be effective.

Consistent with this, LTS tumours demonstrated a broader immune-enriched phenotype, including increased B cell representation, CD8⁺ T cells, and NK cells, indicative of coordinated cytotoxic responses [82]. In contrast, STS tumours showed features of macrophage-driven immunosuppression, consistent with pro-tumorigenic M2 polarisation [83,84]. We also identify GATA6 as a key feature of LTS tumours, consistent with its association with the classical PDAC subtype and immune-enriched microenvironment [42–44]. This supports the idea that tumour-intrinsic transcriptional states shape immune architecture and suggests that GATA6 may serve as a biomarker for patients more likely to benefit from immune-based therapies [43].

An additional layer of insight from this study is the spatial heterogeneity between tumour core and invasive margin TLSs, which revealed distinct immune niches with differential functional implications. While TLSs were present across both regions, their composition, organisation, and association with outcome differed. Invasive margin TLSs were more frequently enriched for myeloid-associated features and exhibited greater variability in structural states, whereas tumour core TLSs in long-term survivors were more often characterised by T cell– and B cell–integrated architectures with higher degrees of organisation and maturity. Importantly, the enrichment of highly organised TLS states (e.g. Cluster 4) at the invasive margin in LTS suggests that this region may act as a critical interface for immune priming and coordination, where immune cells first encounter tumour antigens and initiate structured responses. In contrast, STS tumours displayed a higher prevalence of isolated or poorly integrated TLS configurations across both regions, particularly at the invasive margin, consistent with impaired immune coordination. These findings highlight that TLS function is context-dependent and shaped by tumour geography, with the invasive margin and tumour core representing distinct but complementary immune environments.

Therapeutically, targeting the spatial coordination of TLSs opens new avenues for next-generation tumour treatments. Strategies aimed at inducing TLS formation, enhancing their maturation, or preventing their disruption by immunosuppressive populations could synergise with existing immunotherapies. Zhang et al. showed that the fibroblast chemokine CCL19 promoted B cell recruitment and TLS formation in a colorectal liver metastasis (CRLM) model that prevented tumour growth in mice [85]. Similarly, therapeutic cancer vaccines, including human papillomavirus (HPV) [86] and granulocyte-macrophage colony-stimulating factor (GM-CSF) cytokine secreting tumour vaccines, induced TLSs with proliferating lymphocytes and favourable immune gene signatures, correlating with survival [87]. Biomaterials and synthetic niches such as hydrogels, collagen scaffolds and nanomaterials have shown to induce mature TLSs and enhance ICB efficacy in preclinical models [88,89]. Kuwentari et al. used an injectable hydrogel carrying CXCL13 chemokine and LIGHT cytokine that increased TLS density and maturity, boosted antigen specific T cells, and synergized with anti - PD1 ICB therapy to eradicate melanoma in mice [89]. Progress will rely on defining TLS heterogeneity, developing safe induction strategies (e.g. biomaterials and vaccines), and combining these with ICB and conventional treatments. From a translational perspective, implementation of TLS-based biomarkers in routine clinical practice will depend on assay robustness, reproducibility, and scalability. A pragmatic first step towards clinical translation would be the development of simplified, reproducible assays used to capture key features of TLS presence and maturation. Such approaches could be integrated into existing pathology pipelines and evaluated for inter-observer and inter-laboratory reproducibility with a standardised scoring framework.

## Limitations

Our study has several strengths, including a unique set of treatment naïve PDAC patients and whole-slide spatial analysis, enabling comprehensive assessment of tumour architecture and heterogeneity. However, limitations include the relatively small cohort size, and the non-randomised single-centre design, reflecting the rarity of long-term survivors. A further limitation is the absence of key stromal and vascular populations, from the multiplex panel limiting the interpretation of the stromal and fibrotic microenvironment and its spatial relationship with immune cell populations. Future studies incorporating larger cohorts, spatial transcriptomics, and functional assays will be required to validate and extend these findings.

## Conclusion

In summary, long-term survival in PDAC is driven not by a single immune feature, but by the coordination of transcriptional programmes, cellular organisation, and spatial architecture. LTS tumours exhibit aligned lymphoid signalling, structured TLSs, and coordinated B and T cell organisation, supporting functional anti-tumour immunity. In contrast, the majority of PDAC tumours, represented by the STS cohort, show a breakdown in this coordination, with inflammatory signalling uncoupled from spatial organisation and increased Treg-mediated immunosuppression, resulting in dysfunctional TLSs. Targeting the assembly of functional TLSs by enhancing chemokine signalling, restoring antigen presentation, and reducing immunosuppression represents a promising therapeutic strategy. More broadly, our work defines immune architecture as a tractable target and provides a framework for reprogramming the tumour microenvironment toward coordinated, TLS-driven anti-tumour responses.

## Methods

### Patient cohort

A total of 47 treatment-naïve patients with PDAC who had undergone surgical resection between 2013-2021 from the Royal Surrey NHS Foundation Trust, Guildford, United Kingdom, were included. The study cohort comprised patients stratified by overall survival (OS) and time to disease recurrence. Long-term survivors (LTS) were defined as those with an OS >5 years and were identified retrospectively. Short-term survivors (STS) were defined as patients with an OS of <5 years and were prospectively enrolled. In total, the cohort included 23 LTS and 24 STS cases. The inclusion criteria of this study were as follows: (1) patients with resectable primary PDAC; (2) sufficient tissue size; (3) regions of interest were defined as intact TLSs within the tumour core and invasive margin of PDAC tissues. For quantitative evaluation, we scanned whole slides and regarded the peritumoral region as the range within 1000 µm from the boundary of the tumour nest. The exclusion criteria were as follows: (1) patients who had received additional treatments before surgery; (2) insufficient tissue; (3) unwilling or unable to provide written informed consent; (4) pregnant, or aged <16 years. Formalin-fixed paraffin embedded (FFPE) tissue was retrieved with ethical approval (IRAS project 303613) and a consultant pathologist with a specialist interest in pancreatic diseases identified the TLSs, tumour and margins in all slides.

### Tumour Sampling/Processing

Surgically resected PDAC specimens were fixed in neutral-buffered formalin (NBF) and FFPE sample blocks were prepared. 4µm serial tissue sections were prepared from each block and placed on FLEX immunohistochemistry (IHC)-coated Microscope Slides (DAKO) and the pathologist (IB) marked the regions of interest on a corresponding haematoxylin and eosin (H&E) stained slide.

### Total RNA extraction and NanoString PanCancer IO 360™ gene expression profiling

The tumour region was macro-dissected at the Veterinary Pathology Centre at the University of Surrey. Multiple tissue samples were taken from different depths of the tumour area, in the FFPE tissue blocks. A regular cleaning protocol was applied for processing samples, with the use of ethanol to clean the microtome and metal forceps before sectioning. Five 20μm thick FFPE embedded tumour tissue curls were taken from the PDAC cohort for RNA extraction with meticulous use of RNaseZAP^TM^ (ThermoFisher Scientific, #AM9780) to avoid contamination. RNA was extracted from the curls using the Norgen Biotek FFPE RNA purification kit (Norgen Biotek, Cat. 25300) as per the manufacturer’s instructions. Measurement of the RNA quality and content was performed with the Nanodrop spectrophotometer (ThermoFisher Scientific, #ND-1000) before samples were stored at -80 °C. After extraction, total RNA from FFPE sections were transferred to the Nanostring-facility at University College London for analysis using the nCounter® PanCancer IO 360 Panel Profile panel (Nanostring Technologies, Seattle, WA). Quality check of raw data was conducted using the nSolver v4.0 (Nanostring Technologies) and the nCounter Advanced Analysis software v2.0 (Nanostring Technologies). The housekeeping gene normalization process step for the quantification of gene expression was performed by geNorm algorithm [37][37] in nCounter Advanced Analysis v2.0.115 program (NanoString Technologies, USA). A quality threshold was applied to filter out low-quality or inaccurate samples. As a result, 18 LTS and 16 STS passed the quality check assessments for RNA profiling. Differentially expressed genes (DEGs) between the two selected biological conditions (LTS vs. STS) were analysed with the default option. This panel would allow an understanding of the vital components involved in the complex interplay between the tumour, microenvironment, and immune response in PDAC. The nSolver Advanced Analysis (NanoString IO 360) used predefined gene signatures built into the panel. Gene set pathway scores were generated using the nSolver Advanced Analysis module (NanoString Technologies), which calculates pathway activity scores based on the averaged standardised expression of predefined gene signatures included within the PanCancer IO 360 panel. Gene expression values were standardised (z-score–like scaling) and averaged across all genes in the pathway, with equal weighting and without gene ranking or permutation testing. This approach generated a single numerical pathway score for each gene set in each sample. Clustered gene pathway scores were calculated from normalized expression values of predefined gene sets that were subjected to unsupervised hierarchical clustering to identify similarities in pathway activity across samples.

### Pathway score≈mean standardised expression of genes in the gene set

Cell type scores were generated using the nSolver Advanced Analysis module, which calculates log₂-based scores from the averaged normalized expression of predefined marker genes representing specific immune and stromal cell populations. Scores reflect the relative abundance of each cell type across samples. All genes are weighted equally.

Cell type score≈mean(log₂ normalised expression of marker genes)

### Functional enrichment analysis

Gene Ontology (GO) enrichment and Kyoto Encyclopedia of Genes and Genomes (KEGG) pathway analysis was performed to explore the biological function of DEGs identified from STS vs LTS tumours. The Metascape (http://metascape.org/gp/index.html)[38] database was employed for functional enrichment analysis of the aforementioned list of DEGs, with the results being visually exportable.

### Chromogenic Immunohistochemistry (cIHC)

Molecular subtypes of PDAC were determined by chromogenic immunohistochemistry (cIHC) performed on FFPE tissue sections. The antibody panel and corresponding markers (GATA6, HNF4α, ΔNp63 and KRT14) used for subtype identification can be found in the **Supplementary Materials, Table S1**.

Figure S1 shows the cIHC expression of basal (ΔNp63, KRT14) and classical (HNF4α, GATA6) lineage markers. Tumour regions of interest (ROIs) were first annotated on matched haematoxylin and eosin (H&E)-stained sections and spatially aligned across serial sections stained with individual antibodies. Detection was performed using 3,3’-Diaminobenzidine (DAB) chromogen (brown precipitate), with haematoxylin counterstaining (blue). Serial sections from the same patient specimen were used for each marker, and ×20 magnification using Phenochart (Akoya Biosciences). Quantification of staining intensity for each marker was performed using inForm Analysis Software (Akoya Biosciences) with a machine-learning batch analysis pipeline. Tissue sections (4 µm) were deparaffinised in three 5-min washes of 100% xylene (VWR Chemicals), followed by rehydration through two washes of 100% ethanol (Fisher Scientific). Endogenous peroxidase activity was quenched using 0.3% hydrogen peroxide in methanol (VWR Chemicals) for 20 min, followed by sequential washes in 70% and 50% ethanol and a rinse in deionised water. Antigen retrieval was performed by incubating slides in citrate buffer (pH 6.0, 0.1 M) at boiling temperature for 12 min, followed by a 2-hour cooling period. Immunostaining was conducted using the VECTASTAIN® Elite ABC-HRP Kit (Vector Laboratories). Tissue sections were encircled with a hydrophobic barrier pen, and non-specific antibody binding was blocked with 2.5% horse serum for 15 min in a humidified chamber. Primary antibodies were diluted in phosphate-buffered saline (PBS) containing 1% bovine serum albumin (BSA; Sigma-Aldrich), and 200 µL was applied per slide. Negative controls received PBS/1% BSA only. Slides were incubated overnight at room temperature in a moist chamber. Following PBS washes, slides were incubated with the biotinylated secondary antibody for 30 min at room temperature, washed, and then treated with the ready-to-use avidin–biotin complex (ABC) reagent for 30 min. Colour development was achieved using the 3,3ʹ-diaminobenzidine (DAB) Peroxidase Substrate Kit (Vector Laboratories) for approximately 2 min, followed by rinsing in deionised water to stop the reaction. Nuclear counterstaining was performed with haematoxylin for 1 min, followed by running tap water for 5 min. Slides were dehydrated through graded ethanol and xylene washes, air-dried in a fume hood for 1 hour, and mounted using VectaMount® Express medium with coverslips.

### Fluorescent Multispectral Immunohistochemistry (mIHC)

A nine-colour fluorescent multispectral immunohistochemistry (mIHC) panel was developed and applied to FFPE PDAC tissue sections to characterise the tumour immune microenvironment (TIME) within a single slide. The panel (CD8; CD68; CD4; DCLAMP; CD19; FoxP3; CD20; Pan-CK) was designed to determine the spatial distribution (intratumoural or peritumoural), maturation status, and cellular composition of TLSs within annotated tumour regions and invasive margins. The antigens, corresponding antibodies, Opal™ fluorophores, and their staining order can be found in the **Supplementary Materials, Table S2**.

Manual mIHC was performed using the tyramide signal amplification (TSA) method to detect antigens and enhance signal intensity. TSA amplifies immune-fluorescent (IF) signals through horseradish peroxidase (HRP)–mediated enzymatic activation of tyramide molecules, which covalently bind to tyrosine residues near the target antigen. Following labelling, bound antibodies are removed without disrupting the deposited Opal™ IF signal, thereby allowing sequential target detection with minimal antibody cross-reactivity. Tissue sections (4 µm) were baked overnight at 60 °C, deparaffinised in three 10-min washes of xylene, and rehydrated through graded ethanol (Fisher Scientific) followed by deionised water rinses. Sections were post-fixed in 10% neutral-buffered formalin (Sigma) for 20 min. Antigen retrieval was performed using either citrate-based (AR6, pH 6) or Tris-based (AR9, pH 9) buffers (Akoya Biosciences), selected according to the optimal pH established during chromogenic IHC optimisation. Slides were immersed in the chosen buffer, heated in a microwave at full power for 1 min followed by 15 min at low power, and then cooled for 15 min. After rinsing in deionised water and Tris-buffered saline with 0.05% Tween-20 (TBST), tissue sections were encircled with a hydrophobic barrier pen. Protein blocking solution (200 µL; Akoya) was applied for 10 min and replaced with the appropriately diluted primary antibody. Slides were incubated for 1 hour at room temperature in a humidified chamber, washed three times in TBST (2 min each), and then incubated for 10 min with Opal Polymer HRP secondary antibody (Akoya; anti-mouse and anti-rabbit). Following additional TBST washes, 200 µL of the selected Opal fluorophore diluted in amplification reagent was applied for 10 min, washed, and subjected to antigen stripping using the same heat-induced retrieval method. This staining cycle was repeated sequentially for each antibody.

In the final staining round, AR6 buffer was used prior to nuclear counterstaining with DAPI or Opal 780. After microwave treatment, 200 µL of Opal 780 was applied for 1 h at room temperature, washed, and followed by DAPI staining for 5 min. Slides were washed in TBST and deionised water, then mounted with ProLong®

Diamond Antifade Mountant (Thermo Fisher) and cover slipped. Each staining batch included a negative control, an autofluorescence (AF) control slide, and a DAPI-only slide. Tonsil tissue served as a positive control for all markers. The AF slide was used to visualise intrinsic tissue fluorescence and to distinguish it from Opal-based fluorescent signals.

Optimisation of the Opal™ tyramide signal amplification (TSA)–based multiplex panel involved several key considerations. Spectrally adjacent Opal™ fluorophores were avoided to minimise emission overlap, and each primary antibody underwent optimisation for heat-induced epitope retrieval (HIER), heat stress tolerance, antibody concentration, and fluorophore dilution. The sequential order of antibody application was arranged to prevent co-localisation of markers within the same cellular compartment or cell type across sequential staining cycles.

### Image Acquisition

Whole slide scans of the stained cIHC and mIHC slides were scanned using the PhenoImager™ HT (Akoya Biosciences) at 20 × magnification and saved as a QPTIFF file. For the mIHC slides, regions of interest (ROI) marked by IB were identified and stamped using x20 objective then imported into the InForm Analysis software (Akoya Biosciences) for unmixing and assessment of the stain quality using an algorithm that was created in the inForm® Tissue Analysis software (2.6 version, Akoya Biosciences). Each ROI was spectrally unmixed using a project-specific spectral library created using single-channel dyes and an autofluorescence control. The unmixed images were restitched and analysed in the open-source software package QuPath 0.3.1.

### Cell phenotyping and quantification: Automated cell annotation

For each marker, we aimed to identify and normalize the data relative to a ‘negative’ cell population, characterized by low or negligible marker expression. We employed a two-component GMM (GaussianMixture from scikit-learn, n_components=2) to model the distribution of marker intensities as a mixture of two Gaussian distributions, representing putative ‘positive’ and ‘negative’ cell populations. The GMM estimates the parameters of these two components: the means (μ₁ and μ₂), standard deviations (σ₁ and σ₂), and weights (w₁ and w₂). The weights represent the proportion of cells assigned to each component.

To accurately identify the ‘negative’ cell population, we implemented a robust approach. First, we compared the weights of the two components. If one weight (w_max) was significantly larger (at least four times greater) than the other (w_min), we assigned the component with the larger weight as the ‘negative’ population. This approach prioritises the identification of a potentially large but low-expressing population. Following the identification of the ‘negative’ population (with mean μ_neg and standard deviation σ_neg), we normalised the marker intensities (x) using a Z-score transformation: x(normalised) = (x−μneg)/σneg.

This transformation centres the ‘negative’ population around 0 and scales the data so that the ‘negative’ population has a standard deviation of 1, facilitating consistent comparison across markers and images. After normalisation, we used the SCIMAP package (v2.0.3) in conjunction with the *napari* image viewer to define marker-specific thresholds/gates for cell phenotyping. Specifically, we visualised the normalised marker intensities using *napari* and interactively defined thresholds that effectively separated positive and negative cell populations based on the normalized distributions. These thresholds were then applied across all cells in the dataset using SCIMAP functions.

### Automated TLS identification and TLS Boundary identification

Cells were categorised as ‘TLS’ or ‘non-TLS’ and used as input for the boundary-identification module. “Cell-of-interest” located closer with each were clustered together using custom function based on DBSCAN. ConcaveHull was used to identify the boundary cells. Parameter-selection details are provided in the accompanying Jupyter notebook.

### Cell proportion analysis

Cellular compositions were calculated as percentages of total detected cells per region. To assess differences between patient groups, Multivariate Analysis of Variance (MANOVA) was performed at patient level across phenotypes. Statistical significance (*p*<0.05) is indicated by asterisks in box plots.

### TLS diversity and architectural state analysis

TLS-level features were extracted from a combined dataset of structural, spatial, and cellular metrics. Features with missing values or low variance (≤20 unique values) were removed to ensure robustness. Remaining features were normalised using the bestNormalize package to achieve approximately Gaussian distributions and standardised scaling across features. To improve interpretability and reduce redundancy, features were grouped into biologically meaningful categories as visualised in Supplementary Figure S2, comprising (a) size (e.g. convex area, axis length, cell count); (b) heterogeneity (e.g. intra-TLS variation metrics); (c) complexity (e.g. shape descriptors such as roundness and solidity); (d) organisation (e.g. entropy, lacunarity, Haralick texture features); and (e) cellular composition (B cells, T cells, Tregs, myeloid cells). For each category, a representative score was computed using the first principal component (PCA), capturing the dominant axis of variation. Where appropriate, scores were directionally aligned with biologically interpretable reference features (e.g. size score correlated with cell count). Dimensionality reduction was then performed on the combined feature set using principal component analysis followed by UMAP to project TLSs into a low-dimensional space for clustering. TLSs were clustered using k-means clustering (nstart = 50), with the optimal number of clusters (k = 3–6) selected by minimizing within-cluster variance across all feature scores while enforcing a minimum cluster size (optimal clustering used k=6). Cluster centroids were computed in reduced-dimensional space and mapped back to feature space to characterise each TLS architectural state.

For each cluster, proportions were calculated per group and compared using Fisher’s exact test. Mean feature scores per cluster were also computed to interpret biological differences between TLS states. To assess patient-specific patterns of TLS organisation, we quantified the consistency of cluster assignments within each patient. For each patient, the proportion of TLSs belonging to the dominant cluster was calculated as a measure of within-patient homogeneity. To determine whether TLS architectural states were patient-specific or spatially driven, we performed (a) Adjusted Rand Index (ARI) analysis with permutation testing (10,000 permutations) to assess clustering consistency relative to patient identity; and (b) Chi-squared tests to evaluate association between TLS clusters and patient or anatomical location (tumour vs. invasive regions).

### Higher-order TLS assembly analysis

To quantify TLS architecture beyond cellular composition, we adapted the Tissue Schematics [39] framework to model each TLS as an ordered assembly of local cellular neighbourhoods. Briefly, each TLS was represented as a graph in which nodes corresponded to previously defined neighbourhood classes and edges represented spatial adjacency between neighbourhoods within the same TLS. For clinical interpretability, neighbourhood classes were mapped to biologically meaningful compartments: B-cell zone, B–T zone, dendritic-cell (DC) zone, macrophage zone, non-epithelial zone, T-cell zone, and tumour zone. All downstream analyses were performed at the TLS level and stratified by anatomical context, with invasive (INV) and tumour (TUM) TLSs analysed separately unless otherwise stated.

### Definition of motifs and rules

Within each TLS graph, we identified recurrent local subgraphs, referred to here as motifs [39]. A motif therefore represents a small, recurring spatial building block of TLS organisation, defined by a specific set of neighbourhood classes and their adjacency pattern. Biologically, motifs capture local immune architecture, such as a B-cell/B–T interface or a three-compartment B-cell/B–T/T-cell arrangement.

We then modelled how these local motifs combine into larger structures using assembly rules. In this framework, a rule is not a single neighbourhood pattern, but a transition or relationship between two motifs that co-occur within the same higher-order TLS assembly graph. Thus, motifs describe the local building blocks of TLS structure, whereas rules describe how those building blocks are connected into larger immune architectures. For each motif 𝑚, we represented its topology as a graph and summarised it by its sorted degree sequence, 𝑑(𝑚). For each rule linking a source motif 𝑚_𝑠_to a destination motif 𝑚_𝑑_, the rule family was defined as the ordered pair (𝑑(𝑚_𝑠_), 𝑑(𝑚_𝑑_)). Exact graph transitions were therefore collapsed into a small number of recurring topological rule families, indexed empirically from the observed data. In the present dataset, four major rule families were observed.

### TLS-by-motif and TLS-by-rule matrices

For each TLS 𝑖, we constructed a motif count vector 𝑀_𝑖_ = (𝑀_𝑖1_, …, 𝑀_𝑖𝑝_), where 𝑀_𝑖j_denotes the number of detected instances of motif 𝑗in TLS 𝑖. These motif instances represent motif embeddings within the TLS graph and may overlap; accordingly, motif counts should be interpreted as motif-instance abundance rather than counts of mutually exclusive biological events.

To account for variation in total motif burden across TLSs, motif counts were row-normalised to generate motif proportions:

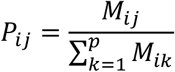

where 𝑃_𝑖j_represents the proportion of all motif instances in TLS 𝑖attributable to motif 𝑗. If no motifs were detected in a TLS, all motif proportions for that TLS were set to zero.

An analogous rule count matrix was constructed for rule families. For each TLS 𝑖and rule family 𝑟, we counted the number of observed motif-to-motif transitions assigned to that rule family, denoted 𝑅_𝑖𝑟_. Rule prevalence was defined as binary presence or absence of the rule family within a TLS:

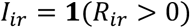

And the within-TLS proportional contribution of a rule family was defined as:

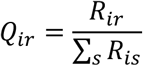

For analyses of “rule proportion when present”, comparisons were restricted to TLSs with 𝑅_𝑖𝑟_ > 0.

### Interpretable annotation of rule families

Because the primary rule-family definition is topological, we generated an additional biological annotation layer to aid clinical interpretation. For each exact motif transition assigned to a given rule family, we took the union of biological compartments represented across the source and destination motifs and collapsed this to a compartment-level interaction class. These interaction classes were then grouped to broader biologically interpretable categories, including pairwise lymphoid interface, multicompartment lymphoid coordination, lymphoid-myeloid interface, tumour-immune interface, and non-epithelial interface. This annotation step was used for interpretation only and did not alter the underlying quantitative rule definitions used for statistical testing.

### High-dimensional megamatrix and Motif module discovery using VDJ-REMIX

To identify spatial patterns and higher-order motif programmes, we analysed the TLS-by-motif proportion matrix using VDJ-REMIX, adapting this framework from adaptive immune receptor repertoire feature analysis to spatial motif composition data. In this setting, each TLS was treated as a sample and each motif proportion as a numeric feature. The analysis was performed within the tumour-region motif space to identify modules of motifs that co-varied across tumour TLSs.

Following the VDJ-REMIX workflow, motif features were filtered to remove low-information variables, and the filtered matrix was used to estimate robust feature-feature correlations. Correlated motifs were then grouped into modules by hierarchical clustering (using hclust package in R) of the correlation matrix. For each module, a single module score was computed as the first principal component (eigengene) of the standardized motif proportion matrix restricted to the motifs belonging to that module. Module scores therefore summarise the dominant pattern of co-occurring motifs within each TLS while preserving interpretability at the feature level. Details of implementation is available at: https://github.com/Bashford-Rogers-lab/vdjremix.

### Rule repertoire similarity

To quantify the reproducibility of higher-order TLS assembly across samples, we compared rule repertoires between TLSs using the Jaccard index. For each TLS 𝑖, we defined its rule repertoire as the set 𝐴_𝑖_of exact rules observed in that TLS. Pairwise similarity between TLSs 𝑖and 𝑗was then calculated as:

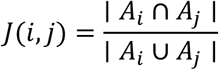

where 𝐽 = 1indicates identical rule repertoires and 𝐽 = 0 indicates no shared rules. Pairwise comparisons were stratified into three categories within each anatomical region: within LTS, within STS, and between survival groups. These comparisons were used to assess whether TLSs from the same clinical group exhibited more concordant rule repertoires than TLSs from different groups.

### Statistical analysis

Neighbourhood composition, motif composition, rule prevalence, rule proportion, module scores, and rule-repertoire similarity were assessed at the TLS level. Rule prevalence between LTS and STS was compared using Fisher’s exact test on binary presence/absence matrices. Rule proportions among TLSs in which the rule was present were compared using two-sided Mann–Whitney U tests. Module scores were compared between survival groups using analysis of variance. For pairwise Jaccard similarity distributions, overall differences across within-LTS, within-STS, and between-group comparisons were assessed using Kruskal–Wallis testing, followed by pairwise Mann–Whitney U tests where appropriate. Multiple testing was controlled using the Benjamini–Hochberg procedure and adjusted 𝑞-values were reported where relevant.

The Kaplan–Meier method was used to plot the survival curves, and the log-rank test was used to compare the differences (GraphPad Prism v9.3.1.). Gene set pathways, logfold2 gene volcano plots and immune cell type deconvolution analysis were performed using the advanced analysis nSolver software tool. Statistical significance was set at *p* value <0.05 for main effects. Normalised NanoString expression data were log2-transformed and analysed for differential expression using the limma package [40]. Log2 fold changes (log2FC) were calculated by comparing mean gene expression between short-term and long-term groups. P-values from moderated t-tests were adjusted using the Benjamini-Hochberg method, and genes with adjusted p < 0.05 and |log2FC| > 1 were considered differentially expressed.

## Authors’ contributions

N.M., S.A., N.E.A., A.E.F., R.B-R conceived and designed the analysis. Computational framework and bioinformatic analyses were developed and executed by S.A. and R.B-R. S.A., S.W., C.V.H., W.G. and R.B-R contributed to code development. S.A and R.B-R. contributed intellectual input/interpretation. I.B., NDK., R.K., R.P.L., T.D.P., E.P., A.R., T.R.W. and A.E.F. provided tumour samples and clinical data. E.V., S.S., T.A.R., D.B., E.G., A.G-W. and M.L.D. provided supervision of the project. N.M., S.A., and R.B-R wrote the paper with inputs from all the authors. All authors contributed to concept and design, read and approved the final manuscript, statistical analysis and interpretation, and critical revision and editing of the manuscript.

## Acknowledgments

This research was funded by The BRIGHT Cancer Care charity, UK (N.M., A.E.F.); Topic of Cancer charity, UK (N.M., N.E.A., A.E.F.); the Royal College of Surgeons of England, UK (N.M., A.E.F.) and the Mason Medical Research Foundation, UK (N.M., A.E.F.). S.A. was funded by Clarendon in partnership with St John’s College. S.W. is supported by Cancer Research UK.

## Ethics approval and consent to participate

All study-related procedures and protocols were in accordance with the ethical guidelines of the Declaration of Helsinki. Patient samples were obtained with Research Ethics Committee (REC) approval (IRAS 303613).

## Code and data availability

All code is available via https://github.com/sakinaamin/pdac_lts_spatial. Processed data and raw images will be available upon reasonable request.

## Supplementary Tables and Figures

**Table S1:** Antibodies used to classify PDAC samples into molecular subtypes: Each PDAC tissue slide was stained with the antibodies listed above to assign the sample to either the squamous/basal or classical subtype.

**Table S2:** Antigens and antibodies selected for use in the human multiplex immunohistochemistry TLS panel.

**Table S3.** Differentially expressed significant genes between LTS and STS tumour regions.

**Table S4.** Transcriptome-derived gene signatures for the detection of tertiary lymphoid structures in human cancers, with identification of 11 key TLS-associated signatures.

**Figure S1:**
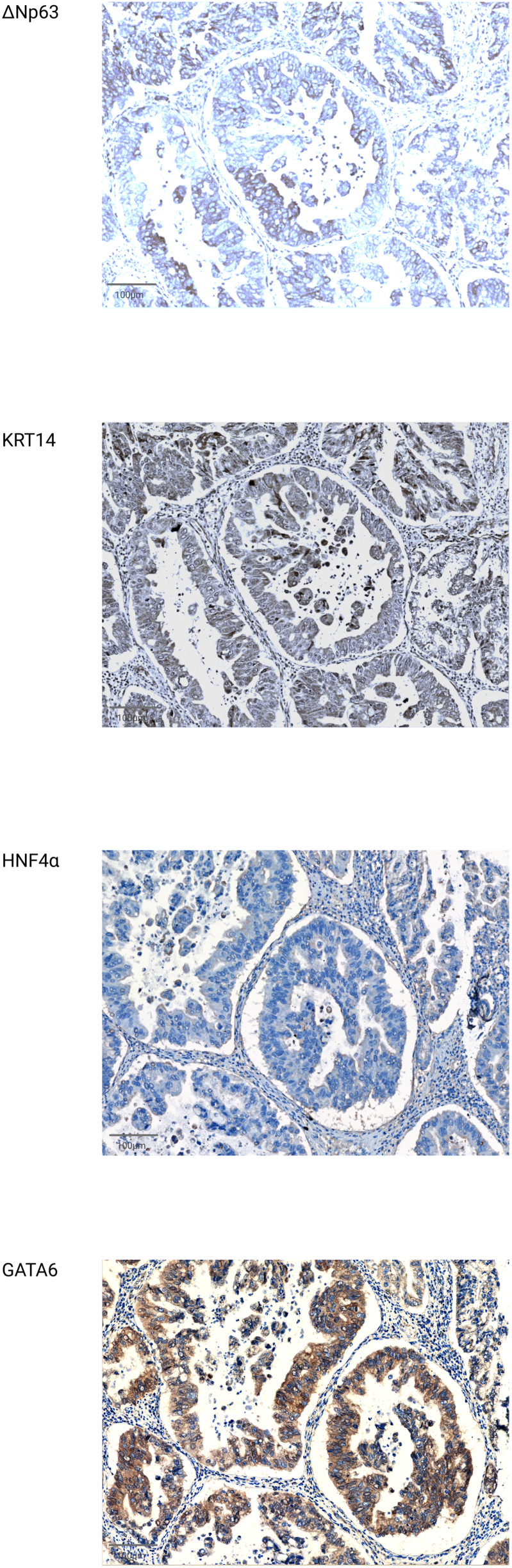
Chromogenic immunohistochemistry reveals basal and classical differentiation states in PDAC tissue sections. Expression of basal (ΔNp63, KRT14) and classical (HNF4α, GATA6) lineage markers. Tumour regions of interest (ROIs) were first annotated on matched haematoxylin and eosin (H&E)-stained sections and spatially aligned across serial sections stained with individual antibodies. Detection was performed using DAB chromogen (brown precipitate), with haematoxylin counterstaining (blue). Representative images show heterogeneous marker expression within the same intratumoral region, highlighting co-existing basal-like and classical differentiation programs. Notably, GATA6 exhibits strong nuclear staining within glandular tumour regions, consistent with a classical differentiation phenotype.

**Figure S2.**
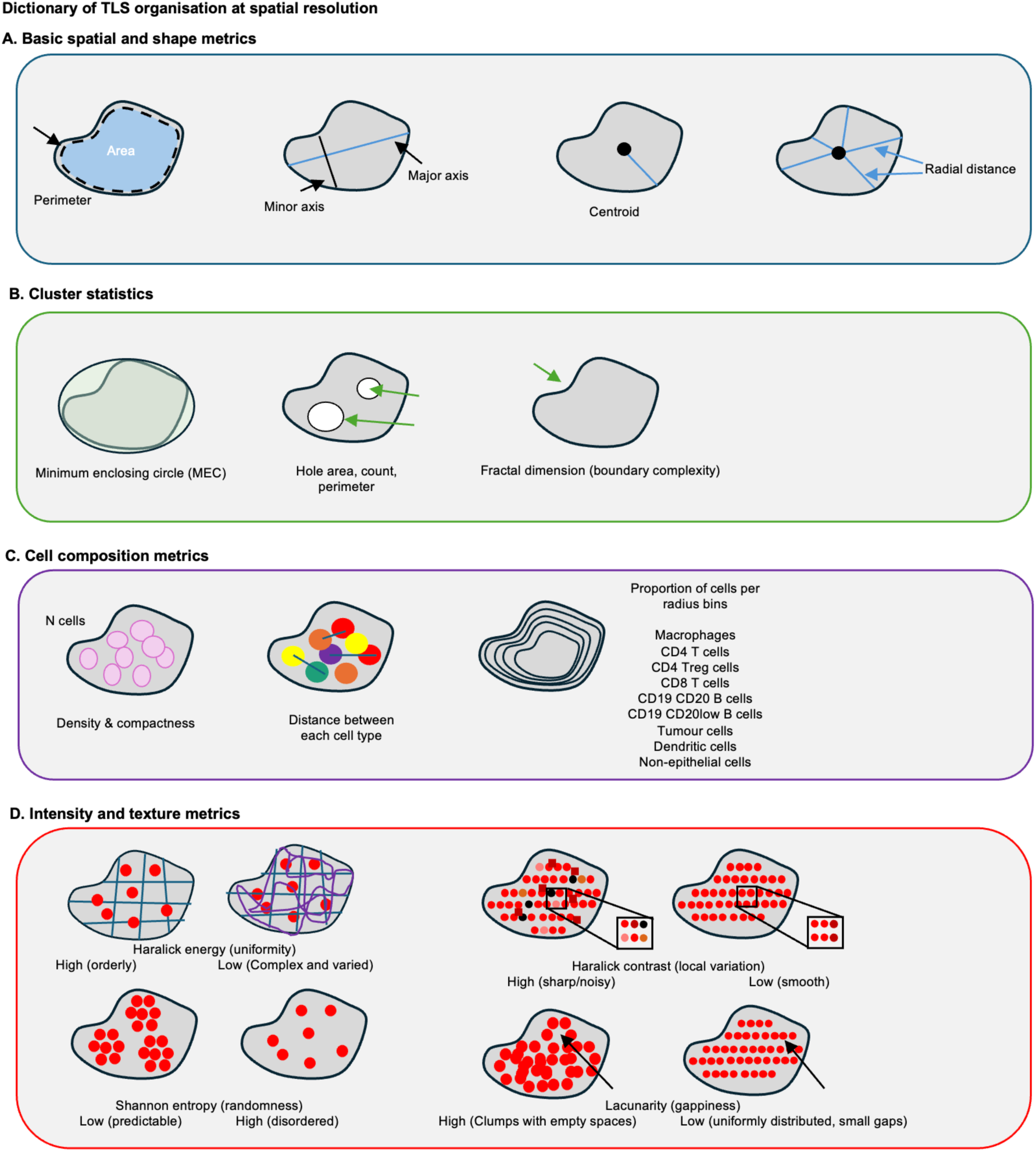
Dictionary of TLS spatial organization and feature extraction metrics. (A) Basic spatial and shape metrics defining fundamental TLS geometry, including area, perimeter, major and minor axes, centroid position, and radial distances. (B) Cluster-level statistics characterizing structural topology, including minimum enclosing circle (MEC), hole-associated features (area, count, perimeter), and fractal dimension as a measure of boundary complexity. (C) Cell composition metrics quantifying TLS internal architecture, including cell density, compactness, inter-cellular distances between distinct lineages, and the radial distribution (spatial bins) of specific immune and non-epithelial cell types. (D) Intensity and texture metrics derived from spatial point patterns to quantify higher-order organization. These include Haralick features (energy for uniformity; contrast for local variation), Shannon entropy for spatial randomness, and lacunarity for characterizing gap distribution and clumping patterns within the TLS.

**Figure S3.**
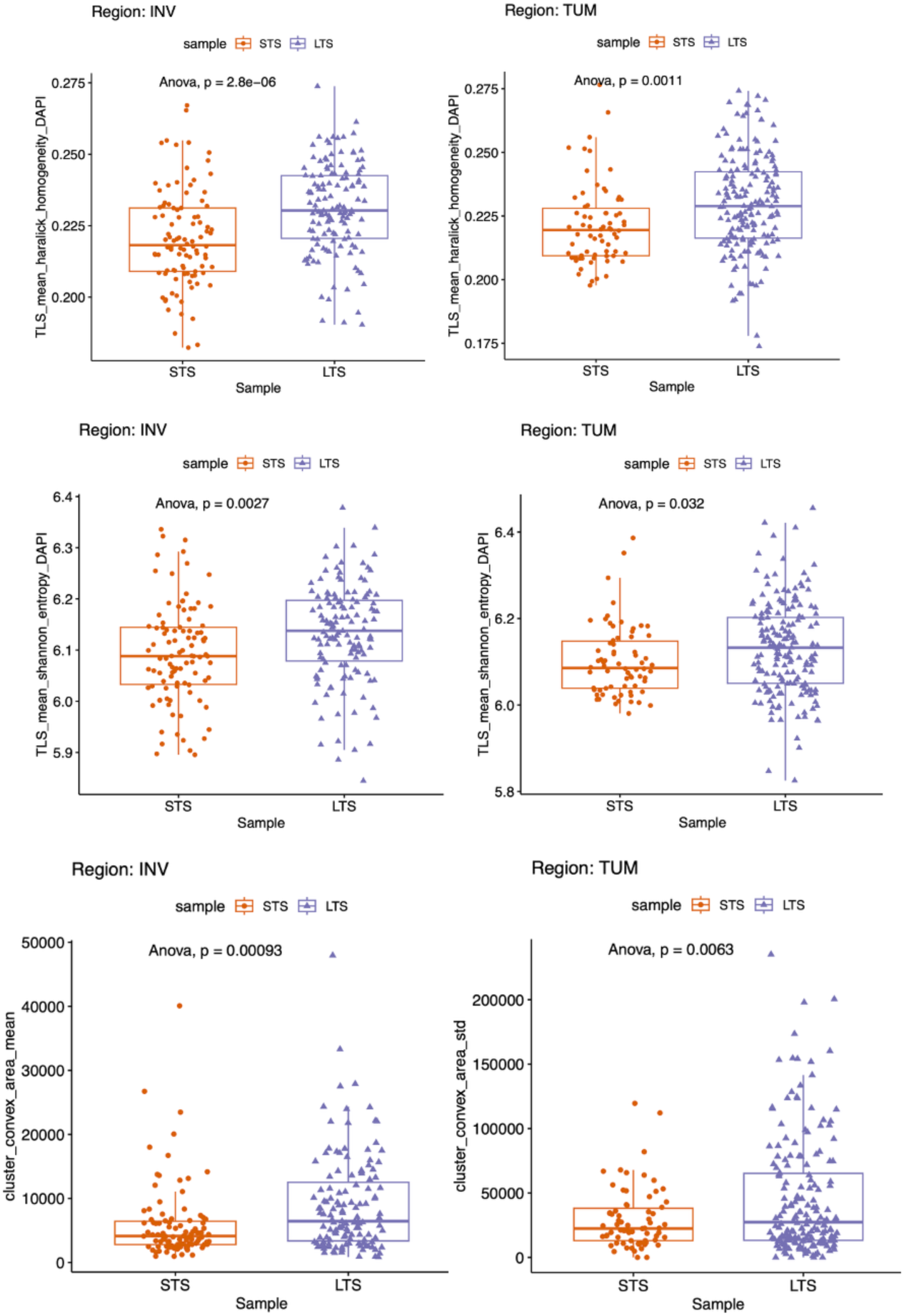
Differential TLS spatial metrics between short- and long-term survivors. Box plots comparing key architectural features of TLSs identified in invasive (INV) and tumour (TUM) regions of short (orange) and long-term survivors (purple).

**Supplementary Figure S4.**
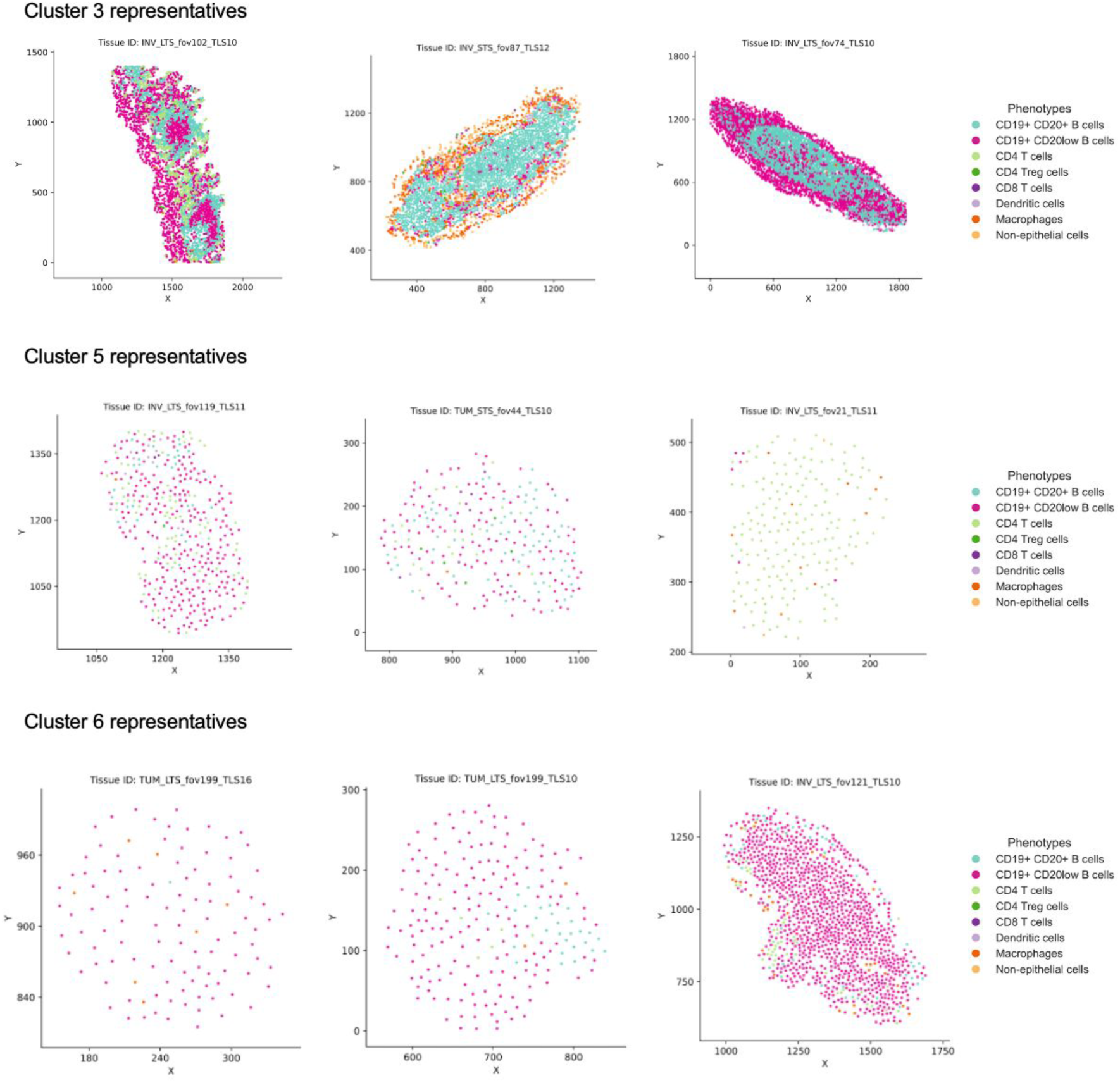
Representative spatial maps illustrating the cellular composition of Cluster 3, 5 and 6. Points represent individual cell phenotypes.

**Figure S5.**
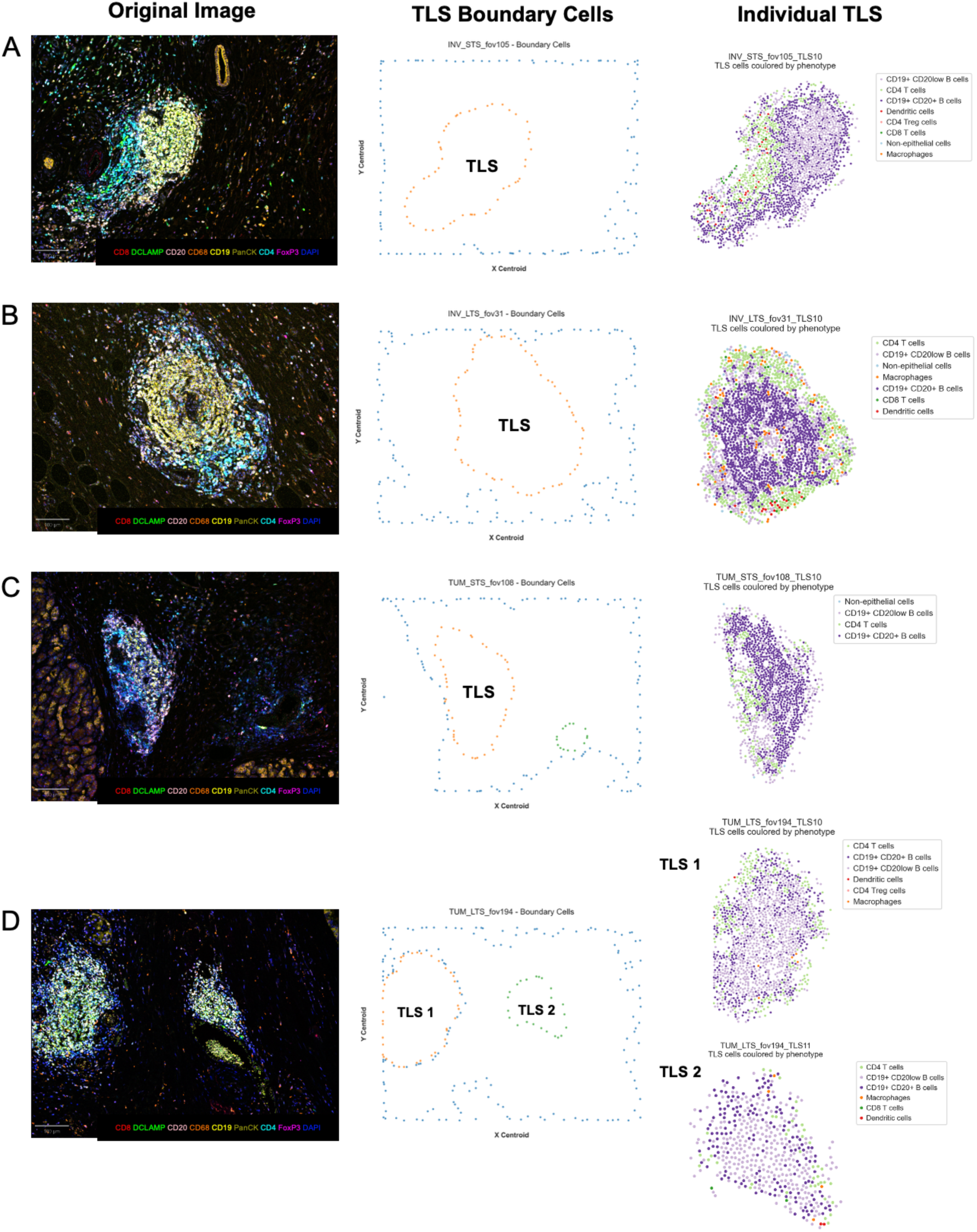
Individual tertiary lymphoid structures (TLS) identification from images across patients and tumour regions. Representative identification of the TLS boundary cells (centre) individual TLS from the original mIHC images (left) across patient groups and regions: A) invasive region of a short-term survivor, B) tumour region of a short-term survivor, C) invasive region of a long-term survivor, and D) tumour region of a long-term survivor. Scale bar at the left bottom side of each image. INV, invasive region; LTS, long-term survivor; STS, short-term survivor; TLS; tertiary lymphoid structure; TUM, tumour region.

**Figure S6.**
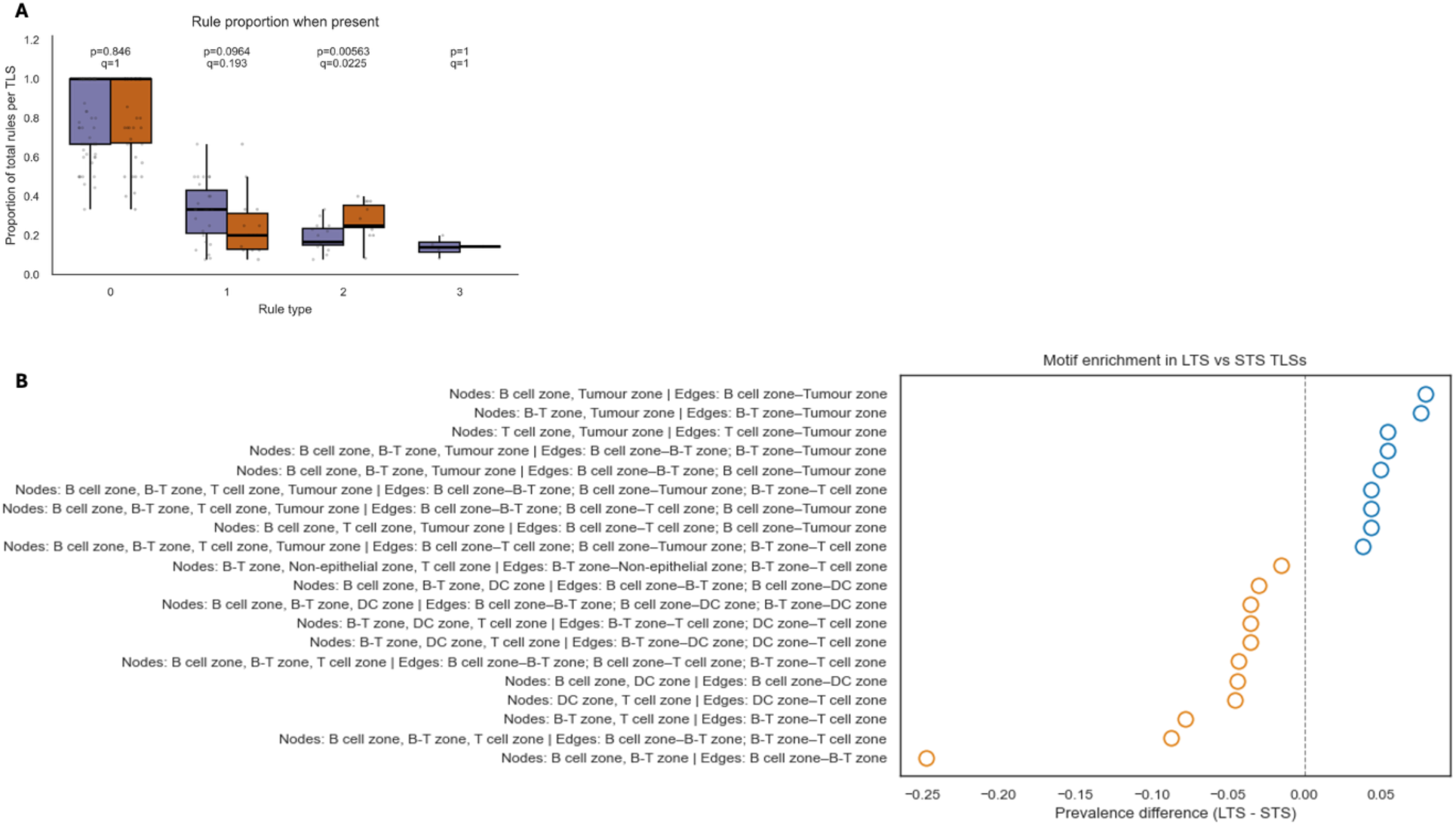
Spatial assembly rules and motif enrichment in LTS versus STS TLSs. (A) Box plots showing the proportion of spatial assembly rules per TLS, categorized by rule type (0–3), in long-term survivors (LTS, purple) and short-term survivors (STS, orange) across all regions. (B) Dot plot illustrating the differential prevalence of specific spatial motifs (defined by nodes and edges of immune zones) between LTS and STS TLSs. Motifs are ranked by prevalence difference (LTS – STS); blue circles indicate motifs enriched in LTS and orange circles indicate motifs enriched in STS.

**Figure S7:**
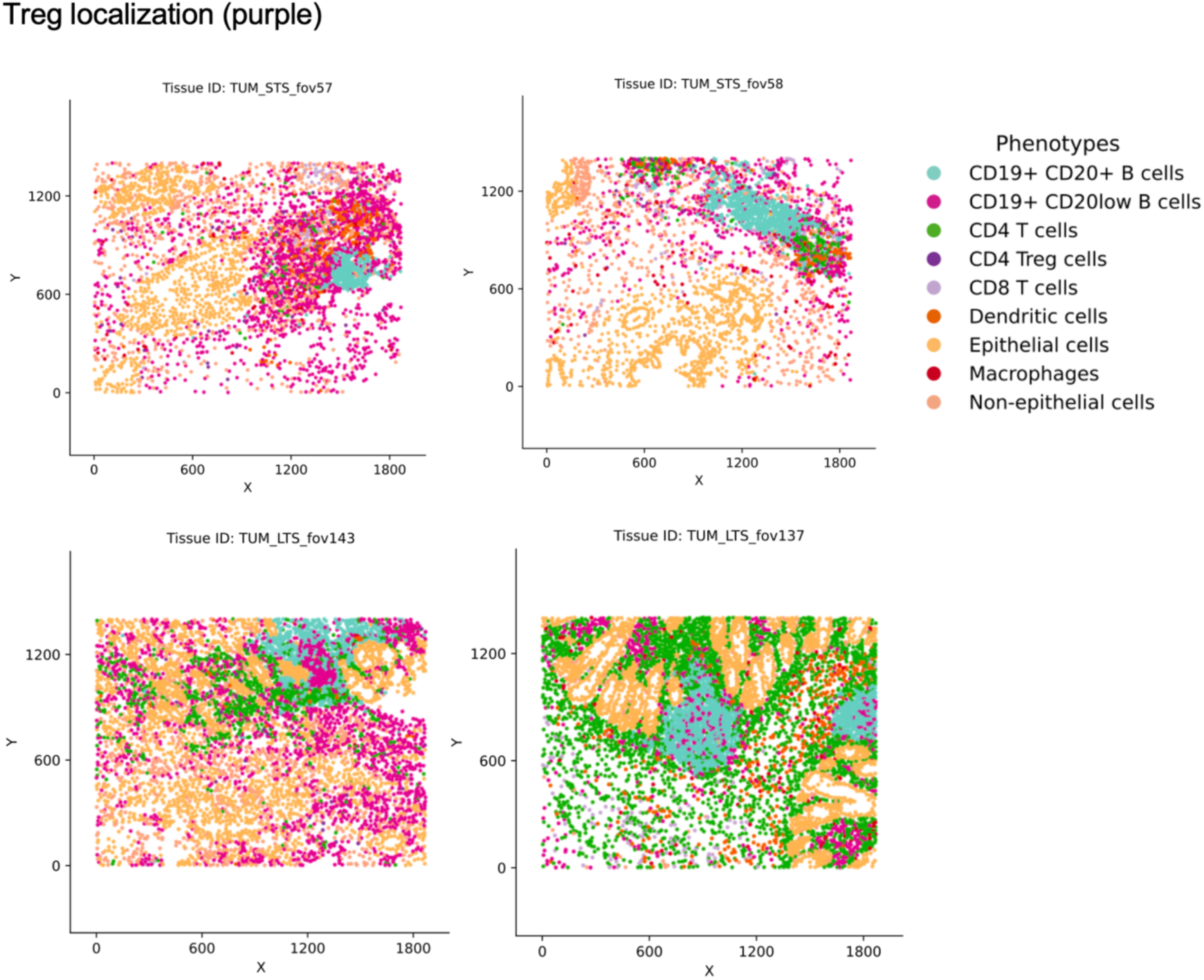
Representative examples of Treg localisation in STS and LTS samples across tumour regions.

**Figure S8:**
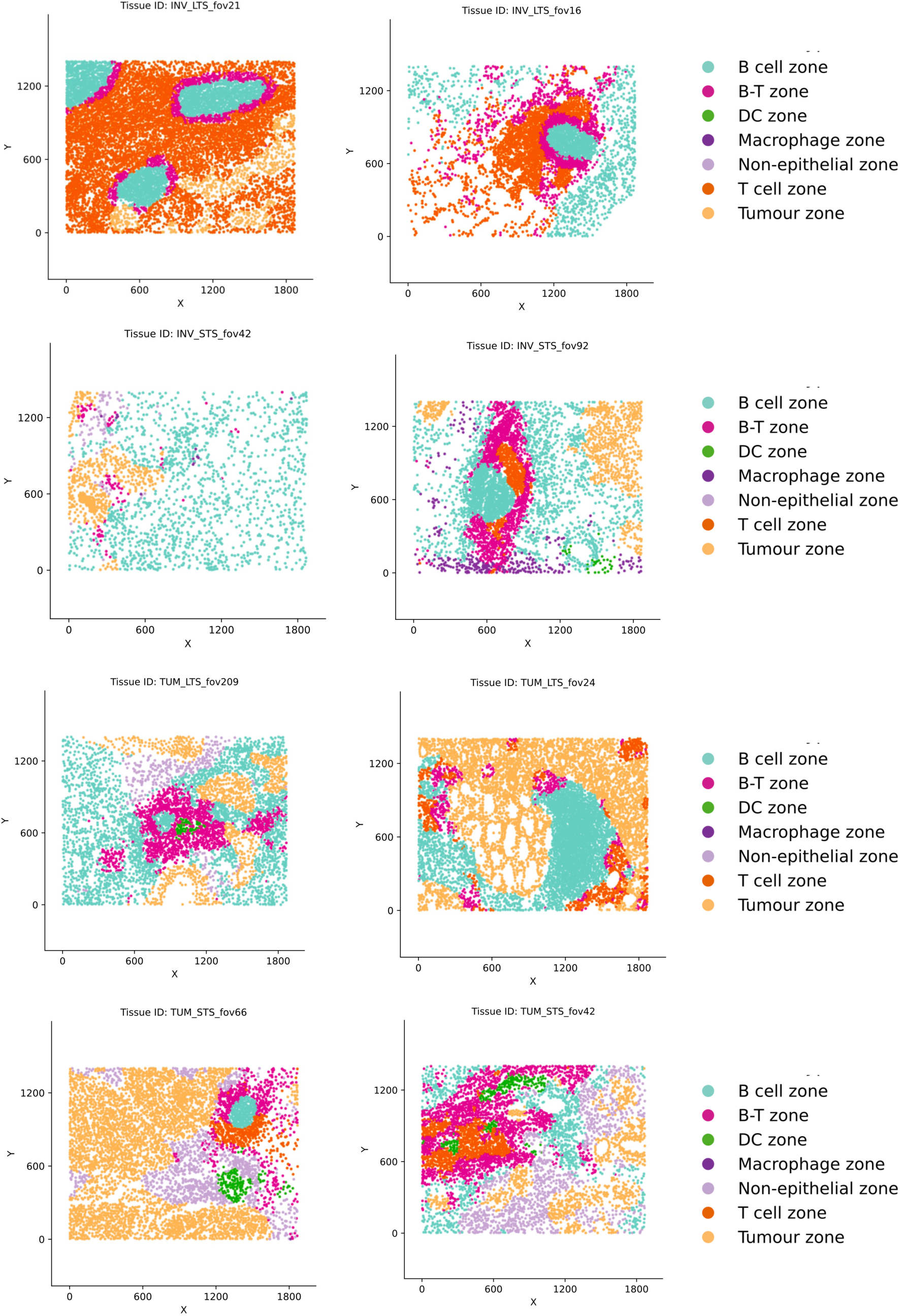
Representative examples of immune zoning in STS and LTS samples across invasive and tumour regions.

**Figure S9.**
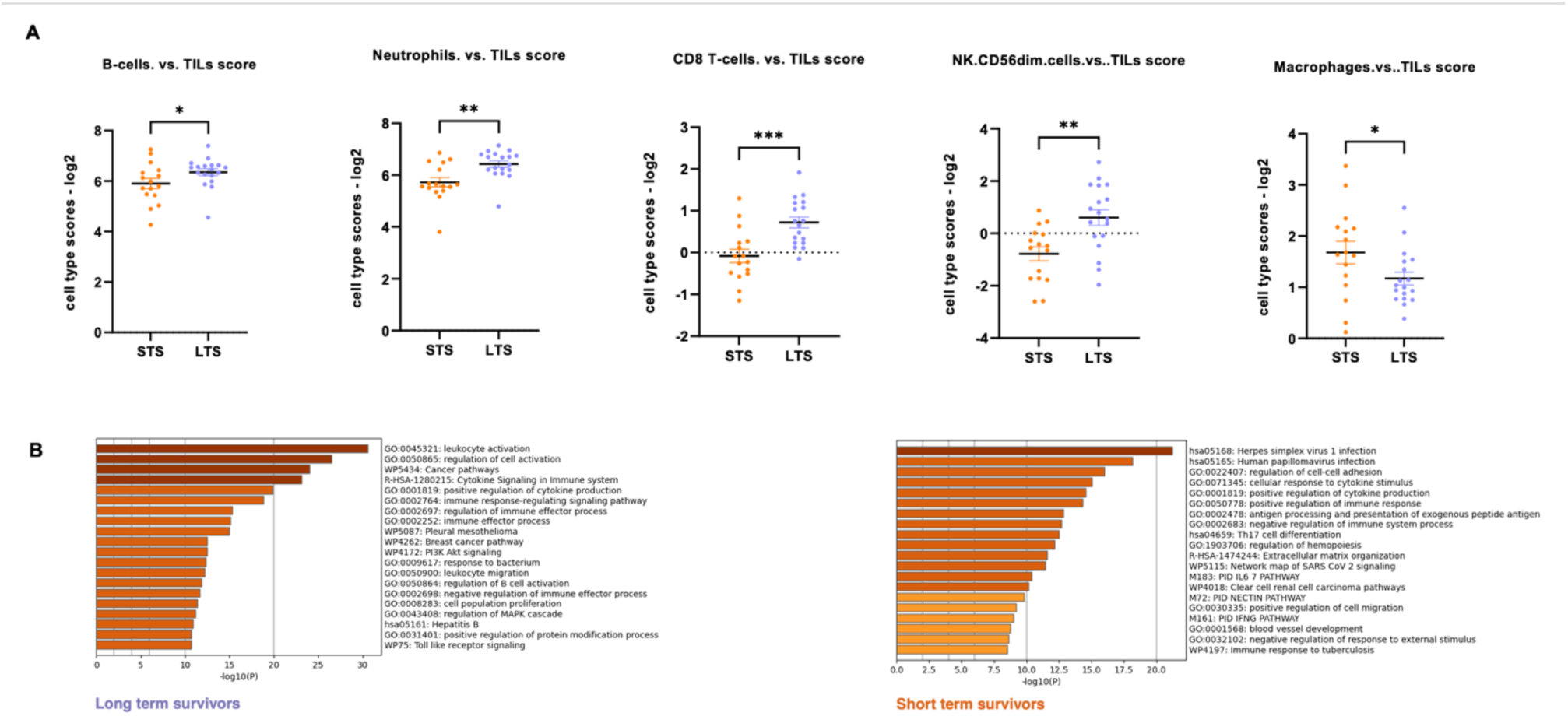
Overview of immune-cell and transcriptomic pathways between long-term and short-term survivors of treatment-naïve PDAC using NanoString PanCancer IO 360™ panel. (A) Significant cell type deconvolutions between LTS (n=18) and STS (n=16). Scatter plot showing immune cell infiltration scores (log₂-scaled relative to total TILs) and data are shown as individual samples with mean ± SEM. Statistical comparison between groups was performed using the Mann–Whitney U test. Significance thresholds: ns = not significant; *P < 0.05; **P < 0.005; ***P < 0.0005. (B) Gene Ontology (GO) and pathway enrichment analysis of differentially expressed genes in LTS (n=18) and STS (n=16). All statistically enriched pathways and hall mark gene sets were identified, and accumulative hypergeometric p-values and enrichment factors were calculated and used for filtering. Pathways shown meet adjusted P < 0.05, with bar lengths representing –log10(adjusted P value).

